# Role of Neural Crest Cells in Establishing Corneal Transparency During Embryonic Development in Mice

**DOI:** 10.64898/2025.12.08.692921

**Authors:** Djida Ghoubay, Cécile Vidal, Quentin Rappeneau, Jasmina Emini, Yorick Gitton, Stéphane Fouquet, Marie-Claire Schanne Klein, Karsten Plaman, Gaël Latour, Vincent Borderie

**Affiliations:** Sorbonne Université, Hôpital National de la Vision des 15-20, INSERM, GRC32, CIC15-20, F-75012 Paris, France; Institut de la Vision, Sorbonne Université, INSERM, CNRS, F-75012 Paris, France; Laboratoire d’Optique et Biosciences, Ecole Polytechnique, CNRS, Inserm, Institut Polytechnique de Paris, F-91128 Palaiseau, France; Laboratoire d’Optique Appliquée, ENSTA, Ecole Polytechnique, CNRS, Institut Polytechnique de Paris, Palaiseau, France; Université Paris-Saclay, 91 190 Gif-Sur-Yvette, France

**Keywords:** Neural crest cells, cornea, corneal transparency, ECM, Sox10, collagen I, SHG, clearing, embryogenesis, embryonic cornea

## Abstract

This study deciphers the exquisitely timed cellular and molecular symphony orchestrating corneal development in mice, revealing how neural crest cells (NCCs) transition from multipotent progenitors to architects of a light-transmitting, mechanically robust stroma. Between embryonic day 10 until birth, NCCs undergo posteroanterior differentiation marked by sequential downregulation of stemness markers, paralleled by collagen I deposition initiating at E12. Stromal expansion and keratocyte-driven ECM remodelling yield precisely aligned collagen fibrils within lamellae a configuration enabling transparency through destructive interference of scattered light. Keratocytes adopt a dendritic morphology to minimize light scatter, while secreting crystallin-like proteins that match refractive indices between cells and matrix. Corneal endothelial maturation and dynamic ECM stratification culminate in a tissue optimized for both optical clarity and structural resilience. Our study elucidates the complex choreography of cellular and molecular events underpinning corneal development, offering novel insights into the acquisition of the cornea’s unique optical and biomechanical properties.

## Introduction

The cornea is the eye’s outermost layer, forming the central part of the ocular surface. It is composed of three layers derived from two embryonic germ tissues: an outer stratified corneal epithelium of surface ectoderm origin, a median stromal layer populated by keratocytes and composed of highly aligned collagen fibrils, and an inner monolayer of endothelial cells covering the posterior corneal surface [1–3]. The latter two layers derive from the neural crests [4–7]. The cornea provides two-thirds of the eye’s optical power and refracts and focuses incident light on the retina [8]. The shape of the anterior corneal surface is essential for maintaining the appropriate corneal refractive power. The corneal viscoelastic biomechanical behavior allows its shape to be maintained despite nychthemeral changes in intraocular pressure and external stress associated, for instance, with eye rubbing. It results from the stromal collagen lamellae micrometric organization [9]. In addition to acting as an external barrier to infectious agents, the cornea is a highly transparent tissue. Corneal transparency is mainly explained by the nanometric organization of the stromal collagen fibrils that run parallel to each other with a constant inter-fibrillar distance [10,11]. This parallel arrangement is thought to result in destructive interference of incoming light rays, thereby reducing scatter and promoting corneal transparency [11]. Keratocytes are a population of mesenchymal neural crest-derived cells sandwiched between the lamellae. They secrete the stromal extracellular matrix (ECM), including collagen fibrils and proteoglycans [12].

During embryonic development, neural crest cells (NCCs) have a sophisticated system for migrating along the nerves to reach their destinations [13]. Prosencephalon and mesencephalon-derived NCC migrate rostrally to form the periocular mesenchyme (POM), contributing to multiple corneal tissues [4,14,15]. These post-migratory NCCs then differentiate into keratocytes, which are responsible for producing and organizing the extracellular matrix (ECM) essential for corneal transparency [16,17]. Various genetic and molecular factors, including transcription factors like Pax6, tightly regulate the differentiation of NCCs into keratocytes and their subsequent activity in ECM production and organization, which is crucial in maintaining neural crest progenitor status and proper eye development [18]. The complete absence of blood and lymphatic vessels characterizes the cornea. This distinctive feature represents an “angiogenic privilege” that naturally prevents vascular infiltration, thereby maintaining the cornea’s exceptional transparency and optical performance [19,20]. Despite significant variations in structure, the cornea retains its transparency across a wide range of species, including non-human primates, humans, fish, and rodents [21].

Abnormal NCC migration, proliferation, or differentiation can result in anterior segment dysgenesis (ASD), a spectrum of ocular malformations involving defects in the cornea, iris, lens, and eyelids [22,23]. Corneal opacity, for instance, ranks as the fifth leading cause of blindness worldwide, affecting approximately 6 million people with moderate to severe visual impairment [24]. Keratoconus, a progressive corneal thinning disorder often beginning in adolescence, demonstrates the dynamic nature of corneal pathology [25]. Despite NCC’s critical role in eye development, our understanding of the transition from opaque corneal precursors to the transparent mature structure remains unclear, in terms of temporal regulation, cell specification, migration and settling, ECM deposition, as well as, topographical orchestration of cell matrix arrangement across corneal domains.

The establishment of corneal transparency, stiffness, and viscoelasticity during embryonic development remains largely unexplored. This gap in our understanding presents a significant opportunity for research in the field of ocular development. Therefore, our study aims to investigate the complex processes that govern the formation of these crucial corneal properties throughout embryogenesis. We elucidate here the cellular interactions that contribute to the acquisition of corneal transparency by tracking neural crest cell migration from embryonic day 10 to postnatal day 0. These developmental stages were selected based on the critical events in corneal development [26]. The development of appropriate mechanical properties by examining collagen synthesis and organization, and the establishment of viscoelastic characteristics of the cornea using complementary imaging modalities such as Full Field Optical Coherence Microscopy (FFOCM), Second Harmonic Generation (SHG) and Transmission Electronic Microscopy (TEM).

## Results

In this study, we analyzed the cellular content in both whole-mount preparations for sample clearing and 3D imaging and cross-sections for confocal microscopy observations of mouse embryos from embryonic day 10 (E10) to postnatal day 0 (P0) to elucidated the mechanism of corneal transparency.

### The dynamic expression patterns of Sox10, Sox9, and HNK1 in the developing cornea

Figure 1 illustrates the whole-mount immunohistochemistry for Sox9, Sox10, and HNK1 at E10. These three markers exhibit distinct expression patterns. Their expression was initiated in the cells as they migrate from the cranial region (Fig 1A, 1B, and 1H). Sox9 was expressed in many cells along the embryo (Fig 1A and 1B; S1 Video). Specifically, Sox9-positive cells were present in several regions, including the midbrain, forebrain and mandibular region (Fig 1A, and 1B). Sox9 expression was also detected in the periocular mesenchyme (POM), the retina, the surface ectoderm, and the forming lens vesicle (Fig 1D). Some Sox9-positive cells were associated with βIII-tubulin, a neuronal marker (Fig 1D).

**Fig 1.**
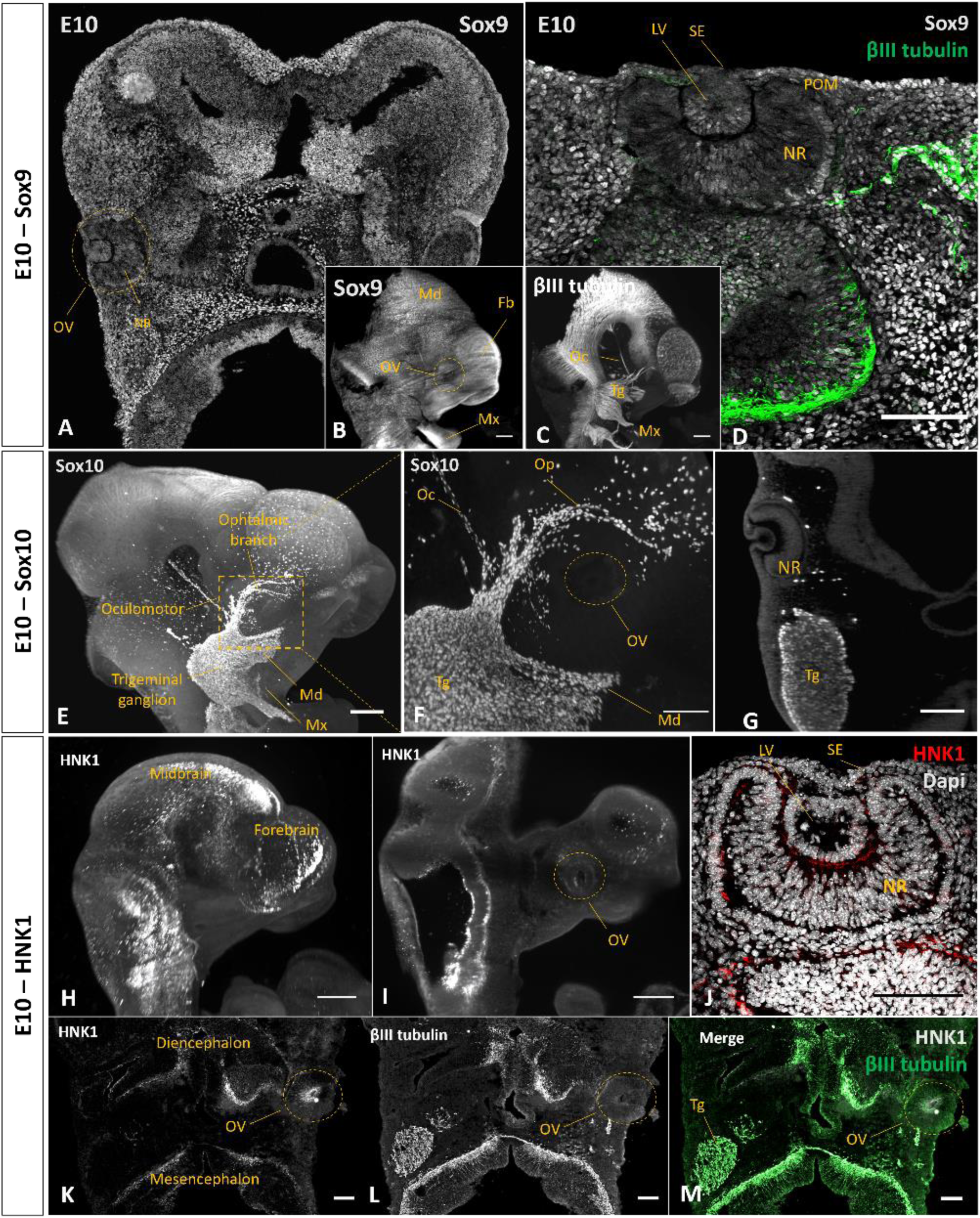
Tracking neural crest cell migration with Sox9, Sox10, and HNK1 markers at E10. (A-D) Sox 9 and BIII tubulin staining in wholemount E10 mouse embryo. (A) Cross-section of the mouse embryo stained with Sox9 in grey. (B) Raw image with Sox 9 positive cells. (C) Raw image with βIII tubulin staining. (D) Cross-section of the optic vesicle region stained with Sox9 and βIII tubulin. (E-G) Sox 10 and βIII tubulin staining in wholemount E10 mouse embryo. (E) Raw image of E10 mouse head with Sox10 positive cells in gray. (F)Raw image of the eye region with Sox10-positive cells (G) Raw images of the optic vesicle section stained with Sox 10. (H-M) HNK1 and BIII tubulin staining in wholemount E10 mouse embryo. (H) Raw image of E10 mouse embryo with HNK1 positive cells in gray. (I) Raw image of E10 mouse head section showing HNK1 positive staining in the cranial region and the OV. (J) Cross-section of the optic vesicle region showing HNK1 staining in the neuroretina. (K-M) The frontal section shows HNK1 and βIII tubulin staining in the head region. L, Lens; LV, Lens vesicle; Ma, Mandibular branch; Mx, Maxillary branch; NR, Neuroretina; Oc, Oculomotor; Op, Ophthalmic branch; OV, Optic vesicle; SE, surface ectoderm; Tg, Trigeminal ganglia. (A, D, F, G, J, K, L, M) Scale bar = 100µm; (B, C, E, H, I) Scale bar = 300µm.

Sox10-positive cells were observed along the trigeminal ganglia and its ophthalmic, maxillary and mandibular branches (Fig 1E and 1F; S2 Video). At this stage, cells present around the optic cup were negative for sox10 marker (Fig1G, S1 Fig). As development progresses, Sox10 expression becomes increasingly restricted to cells associated with developing nerves, particularly those following the ophthalmic and ciliary nerves towards the corneal periphery (Video S3).

HNK-1 is an important carbohydrate epitope that plays a significant role in mouse embryonic development, particularly in neural development and cell migration [27]. HNK-1 was expressed in early migrating NCC. At E10, HNK1 expression was observed in the cranial and trunk region (Fig 1H and 1I; S4 Video). HNK1-positive cells were also observed in the embryonic neuroretina (Fig 1J, 1K).

At E13, Sox9 expression was reduced throughout the embryo, except in the distal regions of developing organs such as the digits, whiskers, intestine, tail tip, and eye (Fig 2A, S5 Video). Sox9-positive cells were observed in the periocular region, and in the surface ectoderm (Fig 3G). The Sox9-expressing cells that persist in the corneal stroma accompany the nerves (Fig 2D-F, S2 Fig). The entry of nerves into the cornea becomes visible from E13 (Fig 2E, 2G), and continue as development progresses (S3 Fig).

**Fig 2.**
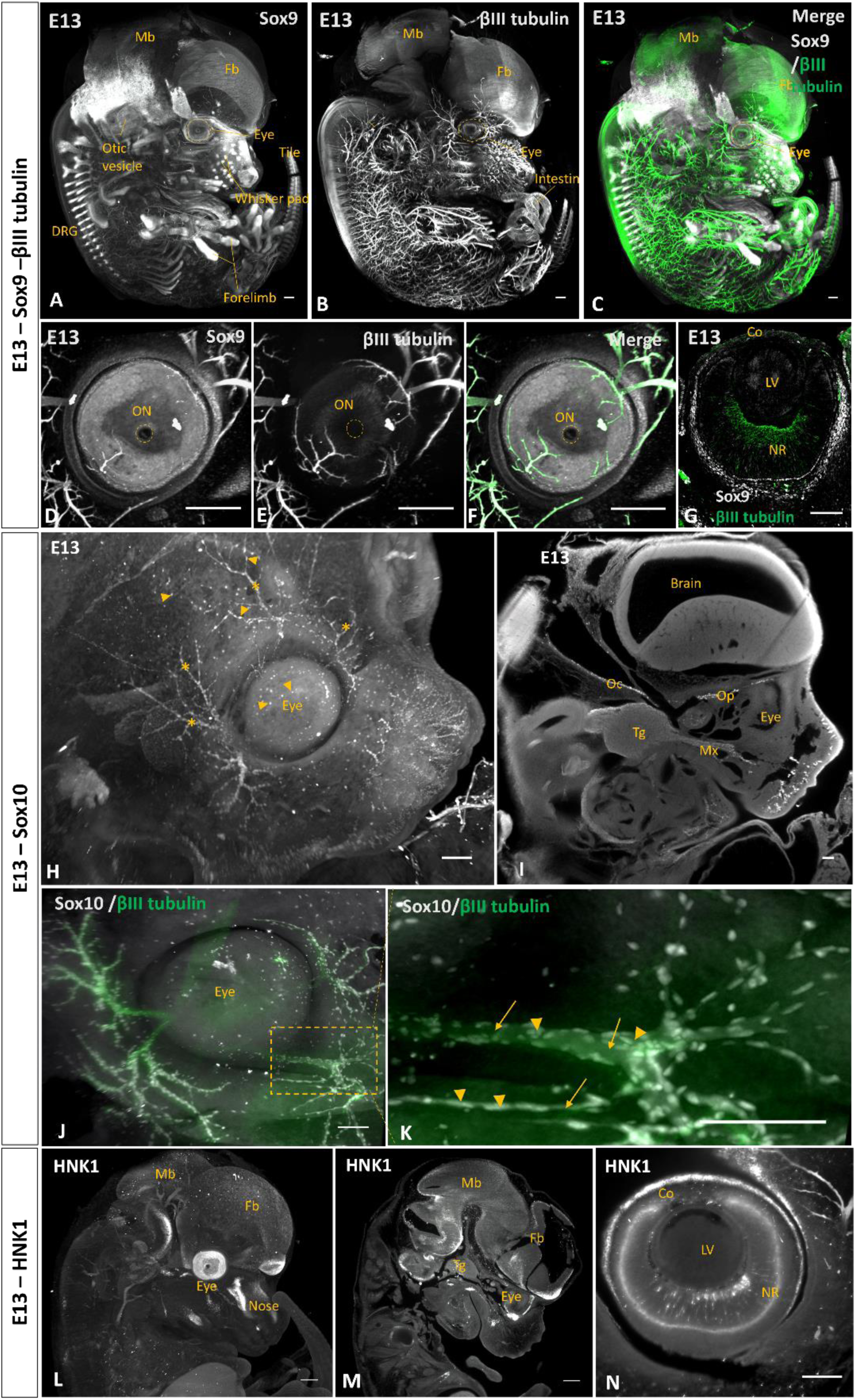
Tracking neural crest cell migration by Sox9, Sox10, and HNK1 markers at E13. (A-G) Sox9 staining in whole-mount E13 mouse embryo. (A) Raw image showing Sox9 staining in gray. (B) Raw image showing βIII tubulin staining in gray. (C) merge of A and B, Sox9 in gray, and βIII tubulin in green. (D-F) The frontal view of the eye at E13 shows Sox 9 (D), βIII tubulin (E), and the merge staining of the two markers (F). (G) Cross-section of the mouse eye at E13 stained with Sox9 in gray and βIII tubulin in green. (H-K) Sox10 staining in E13 mouse embryo. (H) Raw image of E13 mouse head embryo. Sox10-positive cells were present as isolated cells around the eye (arrowheads) and cells close to the nerves (*). (I) Lateral section of E13 head embryo with Sox10 staining. (J) Frontal view of the eye with Sox 10 (gray) and βIII tubulin (green) staining. (K) shows the colocalization of the βIII tubulin in green and Sox 10 positive cells in gray; arrows show the nerves; arrowheads show Sox10 positive cells. (L-N) HNK1 staining in E13 mouse embryo. (L) Raw image of E13 embryo with HNK1 staining. (M) Lateral section of the head E13 embryo with HNK1 localization in the cranial region. (N) Lateral section of the eye region showing HNK1 staining in the neuroretina. Co, Cornea; DRG, Dorsal Root Ganglia; Fb, Forebrain; LV, Lens vesicle; Mb, Midbrain; Mx, Maxillary branch; NR, Neuroretina; Oc, Oculomotor; ON, Optic nerve; Op; Ophthalmic Branch of the trigeminal ganglion. Tg, Trigeminal ganglion. A, B, C, D, E, F, H, I, L, M, N: Scale Bar= 300µm. G, J, K: Scale bar=100µm

**Fig 3.**
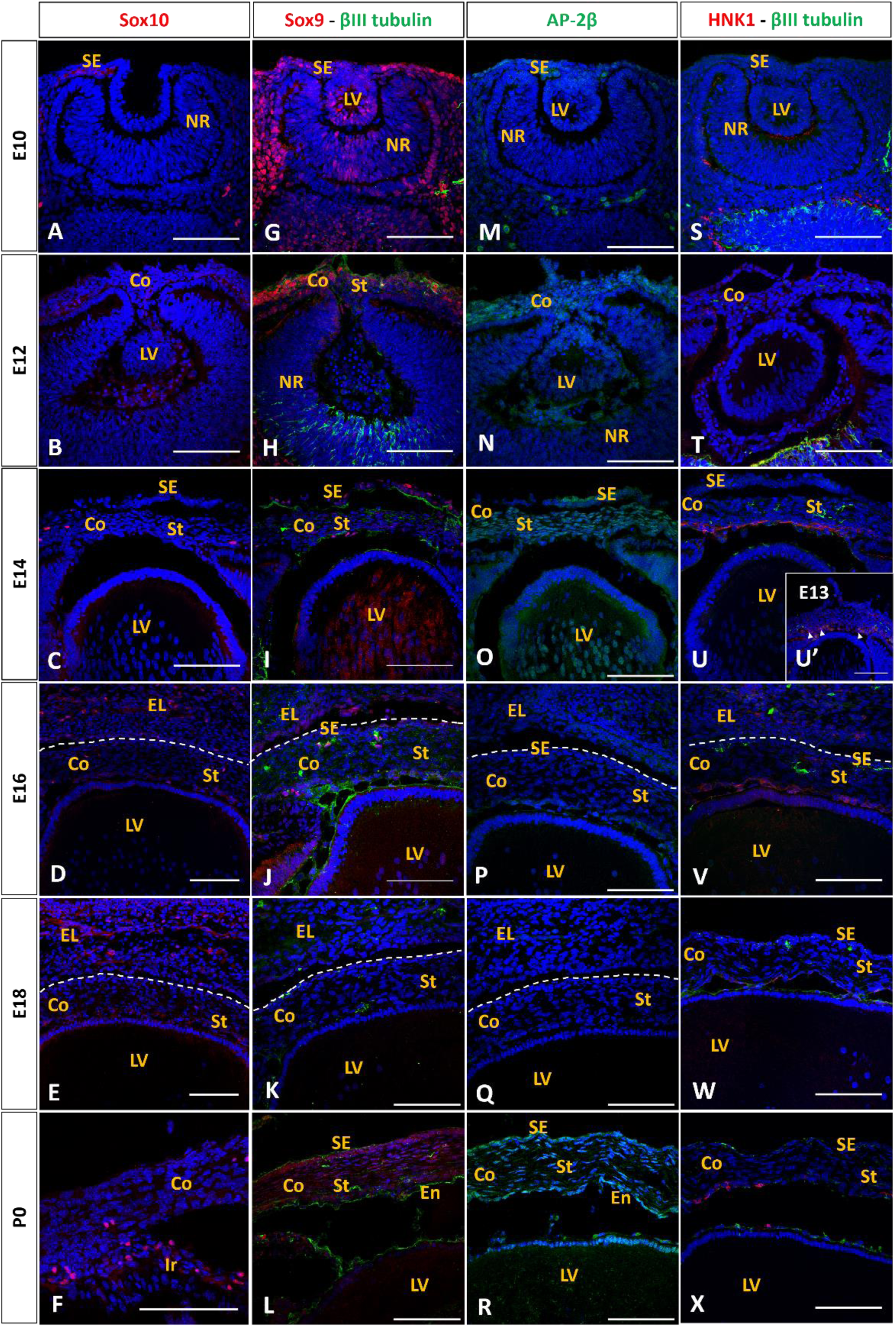
Detailed timeline of neural crest cell migration and differentiation during corneal development in mice. (A-F) Sox10 expression in mouse embryo from E10 to P0 (G-L) Sox 9 and βIII tubulin expression in mouse embryo from E10 to P0 (M-R) AP-2β expression from E10 to P0 in mouse embryo (S-X) HNK1 and βIII tubulin expression from E10 to P0 in mouse embryo. Co, Cornea; En, Endothelium; EL, Eyelid; LV, lens vesicle; NR, neuroretina; SE, surface ectoderm; St, Stroma. Scale bar= 100µm;

Sox10 positive cells were present all around the ocular region and some were also found in the center of the eye (Fig 2H). Figure 2I showed their presence in the trigeminal ganglion, the oculomotor and ophthalmic branch. Sox 10 positive cells surrounding the eye were associated with the nerves as indicated by the double labeling with βIII-tubulin (Fig 2J and 2K). This tight association between Sox10-positive cells and developing nerves continues throughout corneal development (S6 Video).

HNK1 was widely expressed in tissues associated with neural crest cell derivatives (Fig 2L, 2M, and S7 Video). HNK1 was found in different region in the head (Fig 2M) like the follicles of vibrissae, the nasal septum the trigeminal ganglion (S4 Fig). It was also observed in the optic nerve and the ganglion cell (S4 Fig).

Immunostaining of cryopreserved sections revealed the expression of the neural crest markers Sox10, Sox9, HNK1, and AP-2β during ocular development. Sox10 was not detected in the POM (Fig 3A et 3B) but was present in the trigeminal nerve at E10 (S1 Fig) and in a few cells adjacent to the optic vesicle at E12 (S1 Fig). By E14, Sox10-positive cells were localized in the corneal periphery (Fig 3C and S1 Fig), where they colocalized with βIII-tubulin, confirming their association with developing corneal nerves as supported by double-labelling analysis (S6 Video). At P0, Sox10-expressing cells were observed within the iris (Fig 3F).

Sox9 expression was detected at E10 (Fig 3G) and E12 (Fig 3H) within the periocular mesenchyme and surface ectoderm. At E13, Sox9 expression became restricted to a limited population within the retinal pigment epithelium (RPE) and adjacent regions (S2 Fig). Sox9-positive cells were also present in the neuro-retina (S2 Fig). At E15 (S2 Fig) and E16 (Fig 3J), Sox9 expression was observed in the corneal stroma, in cells colocalized with βIII-tubulin.

AP-2β another NCC marker [28], was expressed at E10 in a subset of surface ectodermal cells (Fig 3M) and at E12 in the POM region (Fig 3N). These cells migrated to the space between the lens vesicle and surface ectoderm (S5 Fig), and by subsequent stages, AP-2β-positive cells were observed in the corneal endothelium (S5 Fig at E15). AP-2β expression declined after E16 (Fig 3P) but was reactivated at birth (P0) in all the layers of the cornea (Fig 3R).

In the eye region, HNK1 was observed in the retinal layer at E12 (Fig 3T). It was also observed in a few cells at the periphery of the cornea (Fig 3U’), corresponding to the cells forming the future endothelium as observed at E14 (Fig 3U), where a layer of cells on the posterior side of the cornea was HNK1 positive. HNK1 staining was maintained in the endothelial cells until birth (Fig 3V-3X). Some HNK1-positive cells were observed in adult corneal endothelium (Fig S6A).

### Pax6 expression in the surface ectoderm for proper corneal development

Immunohistochemical analysis showed the expression of the transcription factor Pax6 in the surface ectoderm and the neuroretina at E10 (Fig 4A) and later at E12 when it appeared in the embryonic lens (Fig 4B). It continues to be expressed until birth (Fig 4C-4F). Pax6 was only present in the basal epithelial layer in the adult cornea (Fig S6C). Pax6 staining was absent in the stromal layer at all stage of development (Fig 4A-AF). CK18, an early epithelial differentiation marker in the developing cornea, was present at E10 (Fig 4G). Early expression of CK18 is consistent with its role as one of the first keratins to appear during embryogenesis [29]. CK15, associated with more mature epithelial cells, appeared later at E14, potentially indicating further corneal epithelium differentiation (Fig 4O). Interestingly, CK15 has been proposed as a putative marker of stem cells in various epithelial tissues, including the limbal epithelial stem cells and early transient amplifying cells in the cornea [30]. CK18 and CK15 expression were maintained until birth (Fig 4L and 4R).

**Fig 4.**
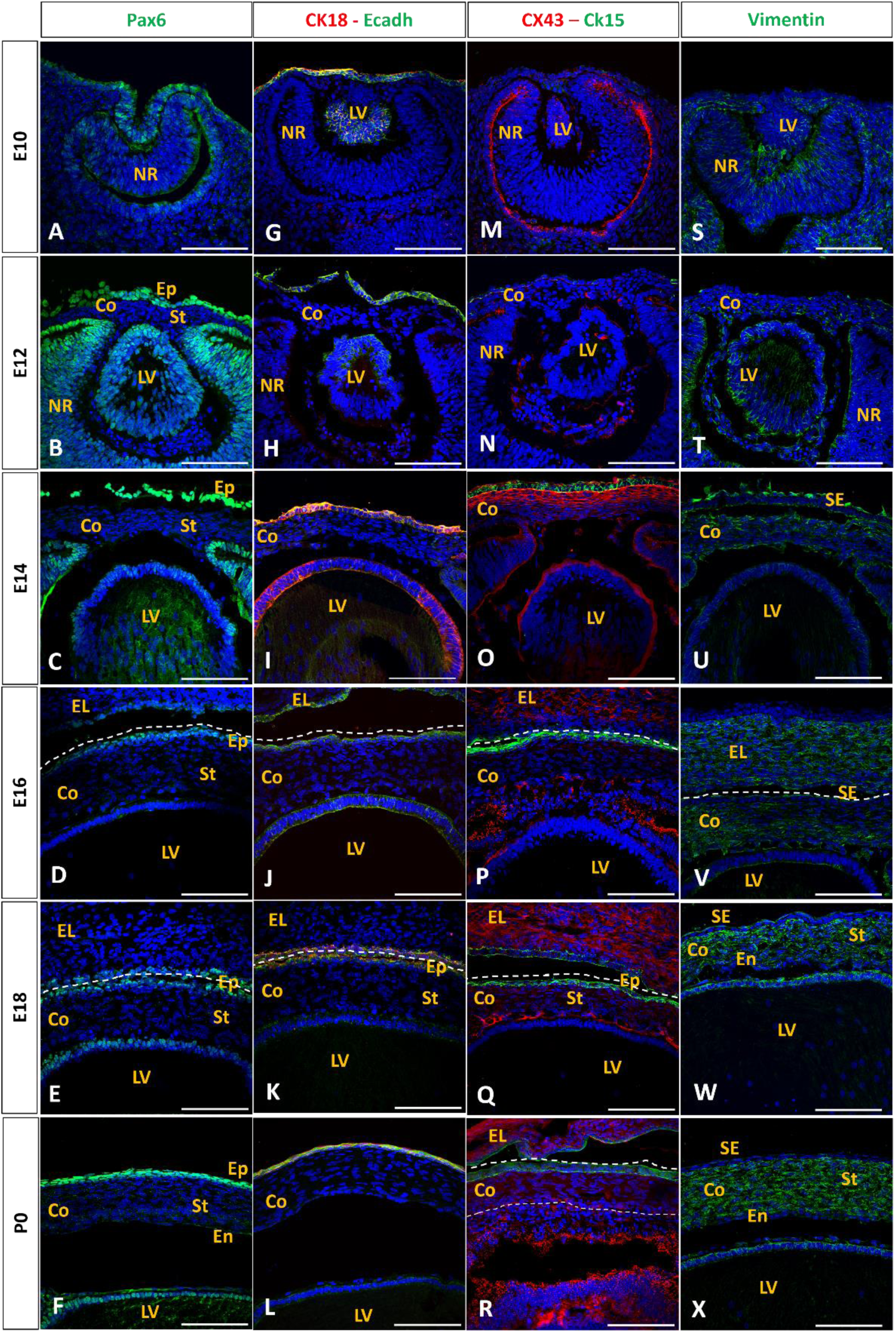
Epithelial cell marker expression and intercellular communication in embryonic mouse cornea. Immunohistochemical analysis of eye vesicle cross-sections from E10 to P0 stained with Pax6, CK18, CK15, Ecadh, CX43, and vimentin. (A-F) Pax6 expression in mouse embryo from E10 to P0. (G-L) CK18 and Ecadh expression in mouse embryo from E10 to P0. (M-R) CX43 and CK15 expression from E10 to P0 in mouse embryo. (S-X) vimentin expression from E10 to P0 in mouse embryo. Co, Cornea; En, Endothelium; LV, lens vesicle; NR, neuroretina; SE, surface ectoderm; St, Stroma. Scale bar=100µm

E-cadherin plays a crucial role in embryonic corneal development in mice [31]. During early eye development, E-cadherin was expressed in the surface ectoderm, which gave rise to the corneal epithelium (Fig 4G). As the lens vesicle separates from the surface ectoderm at E11, E-cadherin expression was maintained in the corneal epithelial cells (Fig 4H), where it functions as a key adhesion molecule [32]. As development progresses, E-cadherin expression in the corneal epithelium is dynamically regulated, with its levels generally decreasing in more differentiated cells [33]. E-cadherin was not expressed in the corneal stroma between E10 and P0 (Fig 4G-4L). During early corneal mouse development, gap junctions are broadly present throughout the corneal stroma as evidenced by immunocytochemistry of Connexin 43 (Cx43) (Fig 4O) playing a crucial role in cell-cell communication [34]. Cx43 expression is maintained until birth in the developing corneal stroma (Fig 4P-4R), and was expressed at E16 in the endothelial layer and continued until birth (Fig 4P-4R). Vimentin is an early marker of cytoplasmic intermediate filaments expressed during embryonic development in mice, particularly in the cornea. Vimentin is found in the periocular mesenchyme at E10 (Fig 4S) and by E12 to P0 in the corneal stroma (Fig 4T-4X). Vimentin seems to be more expressed in the posterior part of the stroma than in the anterior part (Fig 4V). From E18, vimentin is detected in the whole stroma and continue to be expressed in adult cornea (S6E Fig).

### Cell morphology and collagen secretion in the developing embryonic mouse cornea

Trichrome staining of sections reveals the developmental stages from E10 to birth, showing the formation of the surface ectoderm and lens placode invagination from E10 to E11 (Fig 5A). Simultaneously, the NCCs migrate to the space between the lens and surface ectoderm at E11 (Fig 5A). The period between E11 and E14 is characterized by significant stromal expansion. From E13, the cells between the lens and surface ectoderm increase in density (Fig 5A), and notable morphological change occurs from E14 as the stromal cells begin to flatten, in the posterior side of the cornea (Fig 5A, arrows). From E15, the cells were more elongated in both the posterior and the anterior side (Fig 5A, *). By birth (P0), the stroma exhibited a uniform aspect posteriorly with packed collagen than anteriorly where collagen show more space between the lamellae (Fig 5A).

**Fig 5.**
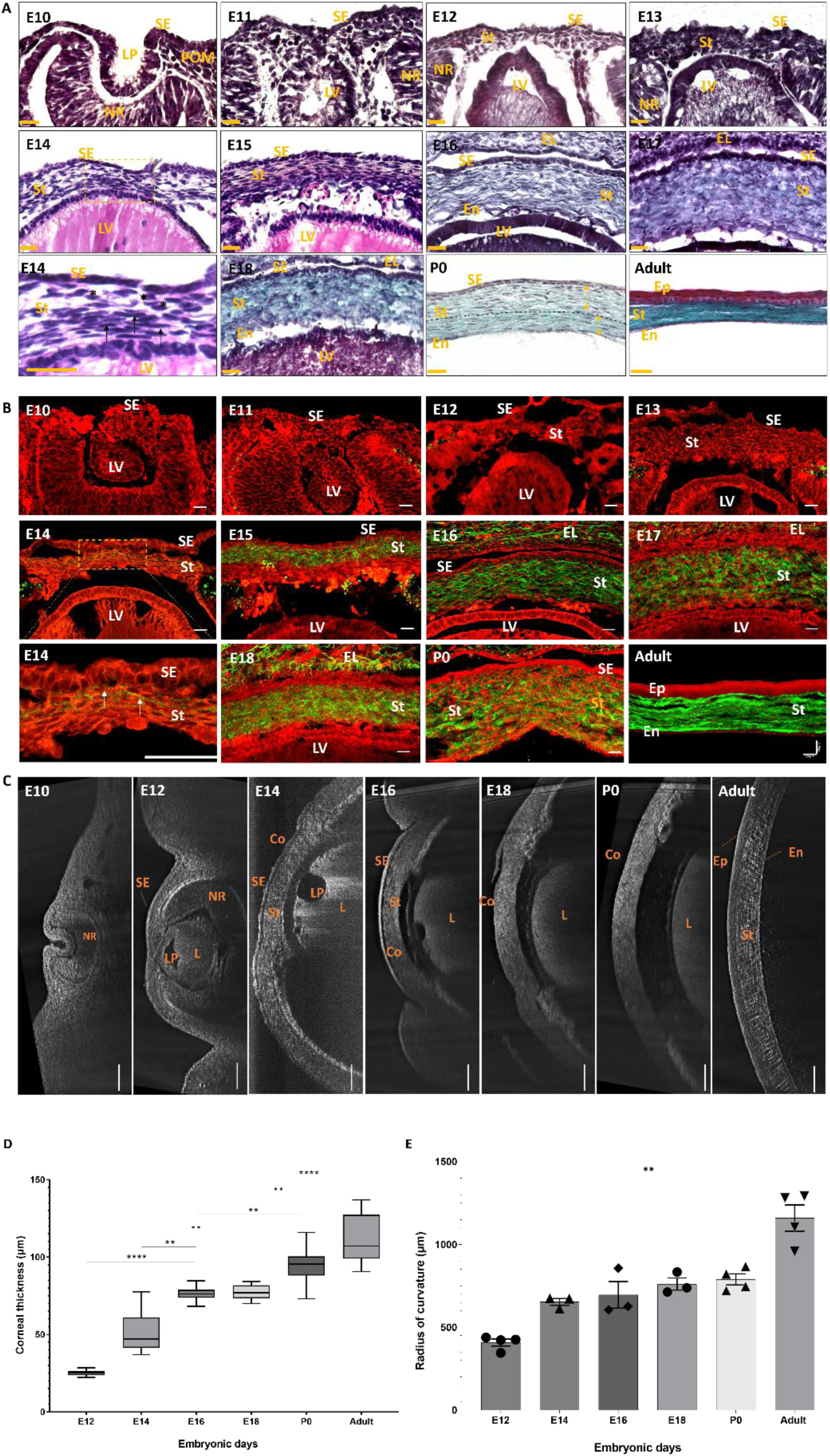
Cell morphology and collagen secretion in the developing embryonic mouse corne. (A) Trichrome staining of cryopreserved mouse embryo sections at different stage of development. nuclei appearing in black, cytoplasm, keratins and muscle fibers in bright red, collagen and mucus in green and blood cells in orange-red. Arrow: flattened cell nucleus, *: rounded cell nucleus. Scale bar = 25µm. (B) Multiphoton images of unlabeled sections. SHG is displayed in green and shows specifically collagen fibrils. 2PEF is displayed in red and shows the cellular morphology of the tissue. SHG is visible from the E14 stage (white arrows). (C) FF-OCT images from E10 to Adult mouse. We observe the corneal development at different stages. (D) Corneal thickness measurement showing the evolution of the cornea at different embryonic days (E) Measurement of the Radius of curvature as the cornea develops. No significant changes in the radius of curvature until birth. Co, Cornea; L, lens; LP: Lens Pit; SE, Surface ectoderm; NR: Neuro-Retina; W, Week. Scale Bar: 100µm; Data are mean ± SEM; ^∗∗^p < 0.01, ^∗∗∗∗^p < 0.0001

SHG imaging revealed the presence of collagen in the corneal stroma as early as E14, visualized as a green signal (Fig 5B). This technique detects individual or groups of few collagen fibrils rather than fully formed lamellae at this early stage. As development progresses, SHG imaging showed an increase in intensity, reflecting collagen fibrils’ gradual organization and densification (Fig 5B). Trichrome staining, while less sensitive, becomes effective in visualizing collagen later in development, typically around E16 (Fig 5A). This method primarily detects collagen lamellae rather than individual fibrils, as it requires a higher density of collagen for visualization.

The embryonic development of the mouse cornea, as observed through lin full-field optical coherence tomography (FF-OCT), revealed a complex structural formation and maturation process. Initially, the cornea begins as a thin layer of surface ectoderm at E12 (Fig 5C). As development progresses, FF-OCT imaging showed a gradual increase in corneal thickness, with the stroma expanding significantly between embryonic days 12 and 16 (Fig 5C and 5D). From E16 to E18, we observed a plateau in the corneal thickness, with no significant changes observed between both stages (Fig 5D). From E18, the thickness of the cornea continued to increase significantly. The corneal radius of curvature increased from 0.4 mm at E12 to 0.79 mm at P0 (Fig 5E). After birth, the radius continued to grow to accommodate the developing eye, reaching an average of 1.16 mm (Fig 5E).

Next, we applied TEM to obtain detailed cellular and structural information, the initiation and intensification of collagen deposition, and the progressive changes in cell morphology and collagen organization throughout development. Collagen deposition initiated around E12 in the sub-epithelial region (Fig 6E), coinciding with the onset of cell proliferation as evidenced by Ki67 immunostaining (Fig 7N). This proliferative activity led to a substantial expansion of the stromal layer, with stromal cells maintaining a rounded shape at this stage (Fig 6F, 6G). During the developmental period from E14 to E16, significant changes occurred in the collagen interfibrillar spacing of the corneal stroma, (Fig 6W-6X). At E14, collagen was predominantly localized in the anterior stroma (Fig 6H), and the interfibrillar spacing measured approximately 57 nm. Between E14 and E16, these spacing decreased, indicating compaction of collagen fibrils (Fig 6W). From E16 to P0, the collagen interfibrillar space in the corneal stroma increased (Fig 6W). This expansion of interfibrillar spacing was likely associated with ongoing collagen deposition and remodelling (Fig 6K, 6N, and 6Q). After birth, however, the interfibrillar spacing decreased, reaching approximately 42 nm (Fig 6T and 6W). In the posterior part of the stroma, collagen fibrils were present at E18 with an interfibrillar spacing of 57 nm, this space decreased as cornea mature reaching 43 nm in the adult cornea (Fig 6W). These results were confirmed by automated calculation of the interfibrillar distance in the posterior part of the cornea (Fig 6Y-6Z).

**Fig 6.**
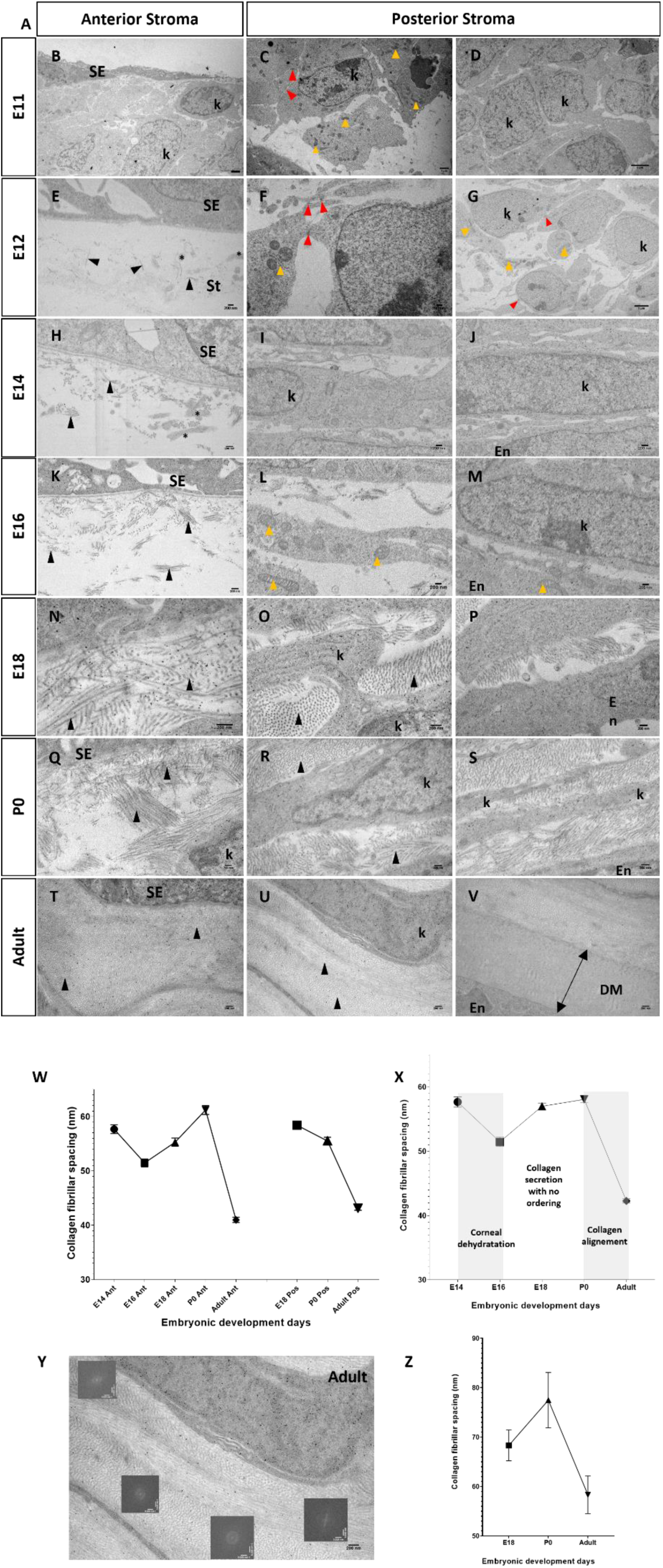
Transmission Electron Microscopy analysis of mouse embryo sections at different stages of development. (B-E) E11; (F-I) E12; (J-M) E14; (N-Q) E16; (R-U) E18; (V-Y) P0. The different images were taken in the anterior stroma, posterior stroma, and endothelium. Black arrowhead, Collagen fibers; Red arrowhead,cell-cell junctions; Yellow arrowhead, mitochondria; Stars, Cell projections. SE, Surface Ectoderm; En, Endothelium; K, Keratocytes. Scal bar: A and C = 1µm; D and G = 2µm; E, F-V = 200nm (W-X) Collagen interfibrillar space analysis in the anterior and posterior stroma with ImageJ. (Y-Z) Collagen interfibrillar space analysis in the whole stromal cornea.

**Figure 7:**
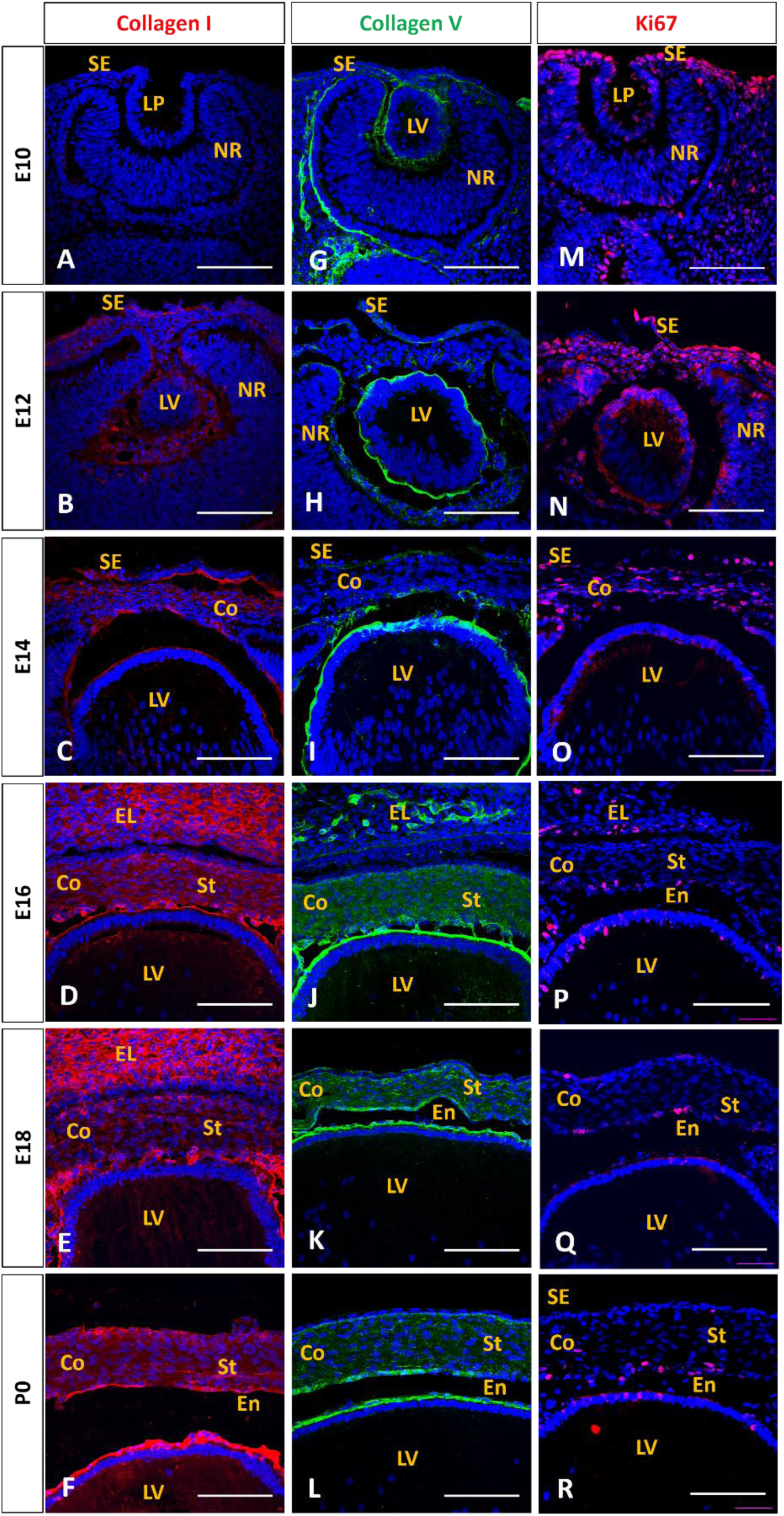
Extracellular matrix expression by corneal stromal cells. Mouse embryonic corneal cross-sections from E10 to P0 immunostained for collagen I (A-F), collagen V (G-L), and Ki67 (M-R). Co, Cornea; EL, Eye Lid; En, Endothelium; LP, Lens Pit; LV, lens vesicle; NR, neuroretina; SE, surface ectoderm; St, Stroma. Scale bar=100µm

Using immunohistochemistry we uncovered distinct patterns of collagen expression, with Collagen I first detected at E12 in the subepithelial region (Fig 7B) and intensifying from E14 to birth (Fig 7B-7F). At the same time, Collagen V showed low levels from E10 to E14 (Fig 7G-7I) and high expression from E16 onwards (Fig 7J-7L). Cells in the corneal stroma all displayed Ki67 staining, suggestive of a massive cell growth from E10 to E14 (Fig 7M-7O). From E16, only cells in the posterior part of the cornea were stained by Ki67, corresponding to the endothelial layer formation (Fig 7P-7R).

## Discussion

Our study provides valuable insights into the migration and differentiation of neural crest cells during corneal development in mouse embryo and the further establishment of corneal physiological properties. The study showed the critical events in corneal development by tracking neural crest cell migration from embryonic day 10 to postnatal day 0. Sox9 is typically the first marker expressed in the developing mouse cornea, appearing in pre-migratory and early migrating neural crest cells [35] as shown in Figure 1. It plays a crucial role in neural crest specification and delamination from the neural tube [36]. As neural crest cells begin to delaminate and migrate, Sox10 expression is initiated (Fig 1), becoming a key transcription factor for their development [37]. Sox10 expression persists during migration and is maintained in glial and melanocyte lineages, though it’s downregulated in many other neural crest derivatives [38,39]. Our data suggest an absence of Sox9 and Sox10 expression as the cornea continues to develop; this is consistent with what was observed by the transcriptomic profile analysis of mouse periocular mesenchyme during the embryonic development of the cornea [40]. At the same time, the transcriptomic profile reveals the expression of S*nai2* and *Twist1* genes in the cornea. Twist1 is involved in craniofacial development and is an inhibitor of Sox9 and Sox10, suggesting a potential role in inhibiting these genes in the cornea [40]. Concurrently, the cell surface carbohydrate epitope HNK1 is expressed on migrating neural crest cells [27]. In mice, HNK1+ cells are considered post-migratory neural crest stem cells, and is often used to identify and isolate NCCs experimentally [41]. Additionally, our findings provide evidence for the neural crest origin of the corneal endothelium, with HNK-1 positive cells forming a monolayer corresponding to the embryonic endothelium by E14, persisting until birth and maintaining expression in some adult endothelial cells. Furthermore, the study documents the migration and differentiation of neural crest cells into corneal stromal cells, with AP-2β expression highlighting its potential role in stromal cell differentiation. AP-2β is highly expressed in the POM and is essential for anterior segment development. Using a mouse model in which AP-2β is conditionally deleted in the NCCs (AP-2β NCC KO), Walker et al. observed structural and phenotypic changes to the stroma and the epithelium associated with AP-2β deletion [28]. The absence of the endothelium characterizes the mutant, and the stroma has a less compact structure, including large gaps leading to corneal opacity [42].

Our results showed that PAX6 expression is continuously maintained in the corneal epithelium during development and in adults’ basal epithelial cell layer [43–45]. PAX6 plays a pivotal regulatory role in embryonic corneal stromal development by orchestrating the interactions between the corneal epithelium and neural crest–derived mesenchymal cells. During early eye formation, PAX6 expression in the surface ectoderm establishes the ocular field and promotes the proper migration and differentiation of cranial neural crest cells that give rise to the corneal stroma [46]. Although PAX6 is not expressed directly in keratocytes, it regulates epithelial-derived signaling pathways—such as FGF, BMP, and TGF-β—that are essential for stromal cell differentiation and extracellular matrix organization [47,48]. Through these mechanisms, PAX6 influences the expression of stromal structural components including collagens type I and V, keratocan, and lumican, which are crucial for the lamellar arrangement and transparency of the cornea [47]. Mutations or haploinsufficiency of PAX6 lead to abnormal stromal development, disrupted collagen organization, and loss of corneal clarity, as observed in aniridia-related keratopathy [49]. Altogether, PAX6 acts as a master transcriptional regulator ensuring coordinated epithelial–mesenchymal signaling, extracellular matrix assembly, and the structural integrity of the developing corneal stroma. Pax6 plays a central role in maintaining corneal avascularity by regulating molecular pathways that inhibit blood vessel formation. Through direct transcriptional control in corneal epithelial cells of the Raver2 and sFlt-1, a VEGF antagonist that sequesters pro-angiogenic VEGF and prevents corneal neovascularization [50].

Our results showed that the cells forming the corneal stroma undergo a crucial process of flattening and collagen secretion, beginning posteriorly and progressing anteriorly (Figure 6). This ordered sequence is fundamental to the cornea’s development and function. It establishes a solid structural foundation and ensures an organized lamellar arrangement essential for transparency [51]. Progressive collagen secretion helps establish and maintain the cornea’s appropriate curvature, which is vital for its optical function. Additionally, this ordered secretion process promotes the differentiation of stromal cells into mature keratocytes and allows for the optimal arrangement of collagen fibers (Figure 4V), further enhancing corneal transparency. Ultimately, this developmental sequence ensures the structured and functional formation of the corneal stroma, which is essential for the optical and biomechanical properties of the mature cornea.

The posteroanterior organization observed in corneal development is a fundamental principle in the embryonic development of numerous other tissues and structures across various species [52,53]. For instance, the body axis formation generally follows a posterior-to-anterior gradient, as seen in the development of the primitive streak in vertebrates [54]. Similarly, the endodermic invagination forming part of the midgut in organisms like Drosophila follows a posteroanterior direction [55]. Even in brain development, although the brain forms at the anterior end of the embryo, its internal regionalization often follows a posteroanterior gradient [56]. This organizational principle is typically controlled by gradients of signaling molecules and sequential gene expression along the embryonic axis, playing a crucial role in establishing the body axis and specifying different body regions.

Intercellular communication is vital for regulating cell growth, proliferation, and differentiation and maintaining tissue homeostasis [34]. In the corneal stroma specifically, gap junctions have been detected at E11 and become more numerous as the cornea matures. Studies have shown that functional gap junctions appear as early as the late 8-cell mouse embryo stage [57]. Overall, gap junctions serve as essential conduits for intercellular signaling during the complex process of corneal stromal development in mouse embryos. The examination of the human keratoconic cornea showed a decrease in the total amount of Cx43 along with a significant downregulation of the active Cx43 isoforms. This may account for a critical mechanism implicated in the keratoconus pathophysiology [34]. The role of gap junctions is identified in regulating the proliferation of neural progenitors and the migration and differentiation of young neurons in the embryonic cerebral cortex [58]. Mutations in the *GJA1* gene, which encodes the gap junction protein Cx43, have been linked to several developmental disorders that affect various body systems [59,60]. These findings highlight the critical role of Cx43 in embryonic development and underscore its importance in the formation and function of ocular structures, craniofacial elements, and other organ systems.

The organization of collagen fibrils in the developing mouse cornea undergoes significant changes during embryonic development, as observed here through imaging. Trichrome staining reveals the progressive deposition and compaction of collagen fibrils in the corneal stroma, with the density and organization increasing as development progresses. Second harmonic generation microscopy provides detailed insight into the three-dimensional arrangement of collagen fibrils because the SHG intensity in each pixel is related to the density and alignment of the fibrils within the focal volume at the micrometer scale, while the SHG pattern in the image shows the distribution of collagen in the field of view at the millimeter scale. It shows a transition from an initially irregular distribution of short elongated structures with low SHG, corresponding to a low-density and disordered network of fibrils with different orientations along the corneal extent, to more organized structures with higher SHG, corresponding to well-aligned and denser assemblies of fibrils aligned along the corneal surface at micrometer and millimeter scales. This figure is likely to be explained by collagen I liquid crystal properties that are evidenced *ex vivo* [61]. TEM offers high-resolution images of the collagen fibril ultrastructure demonstrating their gradual organization into orthogonal bundles. As the cornea develops, collagen fibrils become more aligned and form a complex network of lamellae, which is revealed by the measurement of the distance between the fibrils.

Changes in collagen interfibrillar spacing during corneal development have significant implications for vision development, as they directly affect the cornea’s transparency, mechanical stability, and refractive properties. Transparency is achieved through the uniform diameter and regular spacing of these fibrils, which minimize light scattering and allow light to pass through the cornea with minimal scattering. In the early stages of development, the interfibrillar spacing decreases from day 10 to day 14, coinciding with eye-opening [62]. The compaction of collagen fibrils after birth minimizes light scattering and enhances transparency, critical for transmitting light to the retina [63].

As development progresses, changes in interfibrillar spacing also contribute to the mechanical properties of the cornea. A tightly regulated spacing strengthens the corneal stroma by enhancing fibril density and lamellar cohesion, stabilizing the corneal shape under intraocular pressure. Studies on collagen XII-deficient mice show that decreased interfibrillar spacing leads to increased stiffness but disrupts lamellar organization, highlighting the importance of proper spacing for biomechanical balance [64]. This organization is reported to be closely related to the sophisticated plywood-like organization of the corneal collagen I that represents a remarkable example of biological self-assembly and structural engineering [61,65]. Intrinsic molecular properties, including collagen’s triple-helix structure and specific environmental conditions like pH and ionic strength, drive this unique physicochemical organization. The resulting architecture provides the cornea with exceptional mechanical properties, including remarkable transparency (nearly 90% light transmission), optimal mechanical resistance, and Young’s modulus of approximately 1 MPa [61,65]. The inappropriate deposition of extracellular matrix within the stroma is associated with disorders like keratoconus and progressive loss of corneal clarity [66–68].

The observed plateau in corneal thickness between mouse embryonic days 16 and 18 in our study, while its diameter continues to increase, suggests the onset of a dehydration process during this period. This stabilization in thickness likely indicates a critical developmental phase where the cornea begins to lose excess water, a process essential for its maturation and the development of transparency. The timing of this event aligns with the known activation of the fetal thyroid gland in mice, which typically occurs around mid-gestation (E16.5-E17.5). This correlation supports the hypothesis that thyroid hormone-mediated dehydration plays a crucial role in corneal development, paralleling observations in other species, such as the chick embryo [69]. Exogenous administration of thyroxine before the natural onset of thyroid activity accelerates corneal dehydration and transparency development. Conversely, inhibition of thyroid function by 2-thiouracil delays these changes. Research has demonstrated an inverse linear relationship between corneal hydration levels and light transmission ability, regardless of endocrine treatment [69]. The formation of the corneal endothelium is a critical step in the overall maturation of the cornea, influencing its structure and function. This specialized monolayer regulates corneal hydration through its “pump-leak” mechanism, maintaining the precise water content necessary for stromal organization and transparency. By controlling hydration, the endothelium ensures the regular arrangement of collagen fibrils in the stroma, essential for minimizing light scattering [70]. The corneal endothelium of the mouse, as found in our study, is established at E14 just before the dehydration process initiated by the activation of the fetal thyroid gland.

The mechanisms underlying corneal transparency have been a subject of extensive research and debate. Early studies suggested that transparency is achieved through matching refractive indices of collagen fibrils and the surrounding matrix [71]. Maurice demonstrated convincingly that corneal stroma’s transparency is due to the regular arrangement of collagen fibrils with uniform diameters and distances smaller than the wavelength of visible light [10]. This organization results in destructive interference of scattered light in all but the forward direction, thus permitting transparency. Subsequent studies have expanded on this foundational theory. Hart and Farrell (1969) and Cox *et al* (1970) were the first to show that the regular order within the fibrils does not need to amount to a crystal-like structure. It is in fact sufficient that the fibrils present a local order over distances comparable to the wavelength [72,73]. This order may be mathematically represented by the depth of correlation expressed by the radial distribution (called “pair correlation function” in modern terms), which is closely related to the autocorrelation function. The theory was completed by Benedek (1971), who first took into account the influence of regions devoid of collagen fibrils (sometimes called “lakes”) and Twersky (1975), fully establishing the concept of pair correlation functions applied to cornea [74,75]. Meek and Boote used X-ray scattering to reveal that the collagen fibril organization in the developing cornea is guided by keratocytes, supporting Maurice’s concept of an ordered fibrillar array [76]. Meek and Knupp (2015) confirmed earlier findings by supporting that the cornea’s transparency is maintained as long as the spatial order of fibrils extends over a distance of at least 120 nm [17].

Furthermore, the evolution of corneal stromal collagen organization across vertebrates reveals complexity and structural integrity progression. Koudouna et al. (2018) demonstrate a transition from a simple’plywood-like’ arrangement in fish, characterized by non-interacting orthogonal collagen sheets, to more intricate patterns in higher vertebrates. Amphibians and reptiles show the emergence of broader lamellae with some branching. At the same time, birds exhibit a complex’chicken wire’ pattern with extensive lamellar branching and anastomosing, particularly in the mid-stroma. Mammals diverge from this trend, developing a unique lamellar pattern with increased branching in the anterior stroma, contributing to enhanced mechanical stiffness. This evolutionary progression suggests a correlation between stromal organization complexity and the cornea’s ability to maintain its shape, with higher vertebrates demonstrating more sophisticated structural arrangements that likely confer greater tissue stiffness and shape control [77].

Our study presents a limitation related to the absence of direct measurement of corneal physiological properties. which can reveal critical insights into tissue formation and cell differentiation by linking biomechanical cues to development outcomes. Such measurements are currently highly difficult or impossible in the very small mouse developing-cornea. However, as function is tightly linked to structure and ultrastructure, we could provide new insights into the relationships between cell events and function establishment.

## Conclusion

The intricate timeline of cellular migration, collagen secretion, and matrix organization emerges as a critical determinant in establishing corneal transparency and biomechanical integrity. Our study elucidates the complex choreography of cellular and molecular events underpinning corneal development, offering novel insights into the acquisition of the tissue’s unique optical and biomechanical properties. The sequential interplay between keratocytes, extracellular matrix components, and the hierarchical assembly of collagen fibrils into lamellae reveals a finely tuned process that optimizes light transmission and structural resilience. These findings enhance our understanding of corneal biology and have significant implications for regenerative medicine and tissue engineering approaches aimed at recapitulating the cornea’s sophisticated architecture.

## Acknowledgement

We are grateful to the staff of the Animal and Phenotyping facilities for help. We thank S. Fouquet and Nermine Saidi from the Imaging facility, and the Team S1 from the institute for their help in the clearing and imaging procedures. We thank Alexis Canette (Electron Microscopy Facility, Sorbonne Université, Institut de Biologie Paris-Seine FR3631) for electron microscopy analysis. Thank’s to the IHU FOReSIGHT. This work was supported by grants from Agence Nationale de la Recherche (grant #ANR-21-CE19- 0010-02, CorMecha project). The sponsor or funding organization had no role in the design or conduct of this research.

## Supporting information

**S1 Fig. Sox 10 immunostaining in mouse embryos cross sections**

Immunostaining images of E10, E12, and E14 showing wider views of the optic vesicle, complementing Figure 3, to better visualize the Sox10-positive cells.

Co, Cornea; LP, Lens Pit; LV, lens vesicle; NR, neuroretina; POM, Periocular mesenchyme. Scale bar=100µm

**S2 Fig. Sox9 and βIII tubulin immunostaining in embryos cross sections**

Images from E12 to E15 showing wider views to visualize the co-localization of Sox9 and βIII tubulin in the embryonic eyes.

Co, Cornea; LV, lens vesicle; NR, neuroretina; RPE, Retinal Pigmented Epithelium; RGC, Retinal Ganglion Cell; POM, Periocular mesenchyme. Scale bar=100µm.

**S3 Fig. Innervation process intensifying as development progresses**

Frontal view of the eye at E14, E15 and E17 showing Sox9, βIII tubulin, and the merge staining of the two markers. ON: Optic nerve. Scale bar: 100µm

**S4 Fig. HNK1 staining in different region of the head**

HNK1, is widely expressed in tissues associated with neural crest cell derivatives.

CO, cornea; FV, Follicles of vibrissae; NS, Nasal Septum; ON: Optic nerve; PtC, Parietotectal Cartilage; RGC, Retinal ganglion cell; TG: Trigeminal Ganglion.

Scale bar, A: 500µm, B and C: 100µm.

**S5 Fig. AP2β immunostaining in mouse embryos cross sections**

Images from E12 to E15 showing wider views to visualize AP-2β in the embryonic eyes.

Co, Cornea; LV, lens vesicle; NR, neuroretina; Arrow head show the endothelial layer. Scale bar=100µm.

**S6 Fig. Adult mouse cornea phenotyping**

**(A-F)** Immunostaining of adult mouse cornea cross sections. Some endothelial cells stained with HNK1, nerves by BIII tub (B). Identification of keratocytes with vimentin staining in the stromal layer (E), epithelial stem cells (Pax6 and p63 alpha) in the basal epithelial layer (C and F) and Collagen I in the stromal layer (D). Scale bar: 100µm

**S7 Fig Spectral measurement of interfibrillar distance**

If the cut performed for TEM imaging 842 is perpendicular to the fibril structure, this gives rise to an annular structure of the Fourier spectrum, 843 the radius of which is inversely related to the interfibrillar distance (a). The upper and lower limit for 844 the radius was estimated from the FFT window. If the fibrils are cut with an angle different from 845 90°, these values were determined on the long axis of the elliptical Fourier spectrum (b). If the fibrils 846 lie in the plane of the cut, two distinct spots appear in the Fourier spectrum, which again permits 847 the determination of the upper and lower limit (c). Given that the interfibril distance is inversely 848 related to distances in frequency space, the upper limit obtained for the interfibrillar distance may 849 be slightly overestimated. It is essential to have homogeneous regions where the fibers are clearly 850 distinguishable.

## Methods

### Experimental model and subject details

The Institutional Animal Care and Use Committee approved all animal procedures, which were conducted following relevant guidelines and regulations.

Collection of Mouse Embryos timed pregnancies were established using C57BL/6NJ mice (The Jackson Laboratory). The morning of vaginal plug detection was designated as embryonic day 0 (E0). Embryos were collected at regular intervals from E10, corresponding to the stage of lens vesicle invagination, through to postnatal day 0 (P0). At least three embryos from different litters were collected for each time point. Embryos were dissected free of extraembryonic membranes in ice-cold PBS, imaged for morphological assessment, and then processed immediately for Full-field optical coherence microscopy or fixed in 4% Paraformaldehyde (PFA) in phosphate buffer (0.12M, pH7.4) for further analysis. Reagents references were listed in the key resources table in the supplementary table 1.

### Sample Fixation

All samples were fixed in 4% Paraformaldehyde (PFA) in phosphate buffer at 4°C for 1 to 2 days, depending upon their size. Specimens were rinsed and cold-stored in fresh 1X Phosphate Buffer Saline containing thimerosal (0.1mg/ml).

### Full-field optical coherence microscopy (FFOCM)

The FFOCM instrument used in this study has been described previously [78–80]. Penetration depth depends on tissue content and transparency. Three embryos (6 eyes per time point) for each time from E10 to P0 and adult one, were observed. The acquired images consist of 1024 x 1024 pixels over a 780 x 780 µm en face field of view over a total thickness of 200µm with a 1.6 µm lateral x 1.0 µm axial resolution. One image corresponds to a 1.0 µm-thick corneal section. Images were taken every 1 µm depth step in the central zone of the cornea. 3D image stacks were examined in en face and cross-sectional views using the multiplanar reconstruction (MPR) software provided with the system.

FFOCM corneal cross sections were used to assess corneal thicknesses and radius of curvature measurement. Thicknesses and radius of curvature were measured manually with IMAGEJ® software. Ten equally spaced points across the cross-sections of each cornea (6 per time point), and the mean values were recorded for the measurement of the thickness of the embryonic corneas.

Calculating the corneal radius of curvature using ImageJ begins by opening the corneal image in the software and calibrating the scale. Next, select the circular selection tool from the toolbar and carefully draw a circle that best fits the cornea’s curvature, adjusting its size and position for optimal alignment. Once the circle is fitted correctly, use the’Measure’ function (Analyze > Measure) to obtain the measurements, including the radius. This radius value corresponds to the corneal radius of curvature at the measured location. It’s important to note that this method provides an approximation, as the cornea is not perfectly spherical.

### Line-field confocal optical coherence tomography (LC-OCT)

Line-field confocal OCT is a combination of confocal microscopy and optical coherence tomography [81,82]. This device allows the optimization of the signal in depth. The signal source is related to the spatial variation of the refractive index as in conventional OCT. The acquired images correspond to transverse views (*x*-*z*) of 1200 x 700 µm² with a pixel size of one micrometer. During three-dimensional acquisitions, the transverse scan is performed over 500 µm with a 1 µm step in the y direction. Image processing is performed using IMAGEJ® by summing eight consecutive images to increase the signal-to-noise ratio without blurring the structures. The images obtained here were used for illustration in the figure 5.

### Whole-mount immunolabelling

All antibodies underwent initial testing on cryosections. For whole-mount immunostaining, samples were transferred to a solution containing the primary antibodies diluted in PBSGT. Volume was adapted to cover the sample in total. Next, samples were incubated at 37°C with agitation at 60 rpm for 7 (E10 to E13) to 14 days (E14 to P0). This was followed by six washes of 1hour in PBSGT at RT. Next, secondary antibodies were diluted in PBSGT. Samples were incubated at 37°C in the secondary antibody solution for 3 days and washed six times for 1 hour in PBSGT at RT.

### Agarose Embedding

To facilitate the handling and imaging of immune-labeled samples with the light-sheet microscope, tissue samples were embedded before clearing in 1.5% agarose. Agarose was dissolved in TAE 1X in an Erlenmeyer flask and heated using a microwave (1000W; reaching boiling point) until a homogeneous solution was obtained. Samples were placed in plastic weighing dishes, and cooled agarose (about 45°C) was poured over it, avoiding bubbles. Samples were either totally or partially covered to create a mold. Samples were removed from the mold, and this latter was cleared separately. The agarose was trimmed as close as possible to the sample with a razor blade.

### Sample Clearing

We followed the iDISCO+ protocol [83]. After immunolabelling and embedding, samples were placed in TPP tubes (15 or 50ml) and dehydrated for 1 hour in methanol (20%,40%,60%,80%, and 100% (2x)) under rotating agitation (Stuart, SB3). Methanol volumes were equal to about 5 times the sample volume. The samples were next incubated in a solution of 67% DCM 33% MeOH overnight, followed by 100% DCM for 30 min at RT on a rotator. Lastly, samples were put in 100% DBE for at least 1 day.

### 3D Imaging of specimens

Cleared samples were imaged with a Blaze light-sheet microscope (Miltenyi Biotec) equipped with sCMOS camera 5.5MP (2560×2160 pixels) controlled by Imspector Pro 7.5.3 acquisition software (Miltenyi Biotec). The light-sheet, of a 4μm thickness, was generated by lasers at four different wavelengths (488nm, 561nm, 639nm, 785nm). 1X, 4X or 12X objectives with different magnification lenses 0.6X, 1X and 1.66X were used. Samples were supported by a sample holder from Miltenyi, placed in a tank filled with DBE, and illuminated by the laser light sheet from one or both sides, depending on the size of the samples. Dynamic horizontal focus, which shifts the focus through the sample while imaging, was used to obtain sharper images during acquisition. Mosaics of 3D image tiles were assembled with an overlap of 10% between the tiles. The images were acquired in a 16-bit TIFF format.

### Antibody screening on cryosections

Embryos from E10 to P0 were collected, and the head was dissected for embryos from E15 to P0. Adult eyeballs were dissected. Embryos, heads, and eyes were cryoprotected in a solution of 15% sucrose at 4°C overnight. They were then embedded in a mixture of 7.5% gelatin, 10% sucrose, and 0.12 M phosphate buffer, followed by freezing in isopentane at-55°C and stored at-80°C until needed. Samples were cut into 10µm-thick sections using a cryostat (Leica CM3050) and placed on slides for frozen sections. Gelatin/sucrose was removed by placing the slides in 1× PBS and incubating them at 42°C for 10 min. Cryosections were blocked in PBS-GT (0.2% gelatin, 0.25% Triton-X100 in 1X PBS) for 1 hour and incubated overnight at room temperature (RT) with the primary antibodies diluted in PBSGT (Table S1). After three washes in PBS, species-specific Alexa-conjugated secondary antibodies (Table S1) were incubated in PBS-GT for 1hr at RT. Sections were washed 3 times, 5 min each, with PBST, followed by mounting with fluoromount-G. Imaging was conducted using either an epifluorescence microscope (Leica DM6000) or a confocal microscope (Olympus, FV1000). 12-bit images were processed with ImageJ, where Z-sections were projected on a single plane using the maximum intensity option under the Z-project function. Images were finally converted into RGB color mode, and figures were then assembled.

### Trichrome staining and imaging

This study employed the Masson-Goldner trichrome staining technique on 4 µm thick sections of mouse embryos from E10 to P0. The sections underwent a multi-step staining protocol following thawing and 5-minute rehydration in 37°C PBS. The slides were first stained with Weigert’s hematoxylin for 5 minutes to mark nuclei, followed by rinsing in tap water and distilled water. Sections were then stained with Ponceau Fuchsin for 5 minutes to highlight cytoplasm and muscle fibers. Phosphomolybdic acid was then applied for 1-5 minutes to clear the red staining, immediately followed by 5 minutes of light green staining to visualize collagen. The slides were briefly rinsed in acidified water and then washed in acetic water for 5 minutes. Dehydration was achieved through two baths of absolute alcohol and clarified in three baths of Sub X (xylene substitute). Finally, the slides were mounted using Eukitt. This method allows for differential visualization of tissue structures, with nuclei appearing black, cytoplasm and muscle fibers bright red, and collagen green, thus providing a detailed analysis of murine embryonic tissue morphology and composition at various developmental stages.

Light microscope images of stained sections were scanned with the Hamamatsu 174 NanoZoomer Digital Pathology (NDP) 2.0 HT (Hamamatsu Photonics, Massy, France).

### Scanning Electron Microscopy Analysis

Corneas from E10 to P0 and from adult mice (2 for each time) were dissected and fixed via careful immersion in a solution of 2.5% (v/v) glutaraldehyde in 0.1 M sodium cacodylate buffer pH 7.2 at room temperature. They were then washed three times for 5 min with 0.1 M sodium cacodylate buffer and dehydrated with increasing ethanol concentrations at room temperature (50– 70– 90%- 3 × 100%, v/v with ultra-pure water) for 10 min at each step. After immersion in acetone and epoxy resin, samples were placed in flat molds (allowing positioning for precise transversal sectioning), and polymerization was performed by incubation for 72 h at 60 °C. Ultrathin sections of 70 nm were cut with a UCT ultramicrotome (Ultracut, Leica microsystems) and deposited on 200 mesh copper grids (Electron Microscopy Sciences). Observations were performed using a LaB_6_ JEM 2100 HC transmission electron microscope (Jeol, Croissy sur seine, France) at 120 kV and with an objective aperture adjusted for each sample and magnification.

### Collagen interfibrillar space measurement

A systematic method was employed at various embryonic and postnatal development stages to analyze the interfibrillar distance in the corneal stroma. The study covered stages E14, E16, E17, E18, P0, and adult. The interfibrillar space was analyzed using ImageJ’s Fast Fourier Transform (FFT) function. Regions of interest (ROI) were defined in ImageJ on TEM images of corneal cross-sections within homogeneous regions containing distinguishable collagen fibril structures in the anterior or the posterior part of the stromal layer. If the cut performed for TEM imaging is perpendicular to the fibril structure, this gives rise to an annular structure of the Fourier spectrum, the radius of which is inversely related to the interfibrillar distance (S7A Fig). The upper and lower limit for the radius were estimated from the FFT window. If the fibrils are cut with an angle different from 90°, these values were determined on the long axis of the elliptical Fourier spectrum (S7B Fig). If the fibrils lie in the plane of the cut, two distinct spots appear in the Fourier spectrum, which again permits the determination of the upper and lower limit (S7C Fig). Given that the interfibril distance is inversely related to distances in frequency space, the upper limit obtained for the interfibrillar distance may be slightly overestimated. It is essential to have homogeneous regions where the fibers are clearly distinguishable. The cuts are not made in a plane perpendicular at all stages. Therefore, only a few images could be analyzed using the Fourier method, making it necessary to use the manual method in order to obtain statistically representative analyses.

ImageJ software was used to measure more than 100 interfibrillar distances in each image manually. Perpendicular line was drawn between two adjacent fibers, and the distance between them was calculated. At each stage, images were taken from the stroma’s anterior and posterior sides. Three images per side were analyzed, totaling six images per developmental stage. At stages E14 and E16, collagen fibers were observed only on the anterior side. From E17 onwards through to the adult stage, collagen fibers were present on both the anterior and posterior sides of the stroma.

### Multiphoton microscopy

Multiphoton microscopy was performed using a custom-developed laser scanning upright microscope to study the characteristics of the extracellular matrix in the developed animal model, as previously described [84,85]. E10 to P0 Embryos were used for this experiment without any labeling. The presence of collagen fibrils was assessed by second harmonic generation (SHG), which is a multiphoton mode of contrast highly specific for unlabeled fibrillar collagen, with unequaled sensitivity [86,87]. The SHG intensity depends on the density of the collagen fibrils and their organization within the focal volume: large SHG is obtained for dense assemblies of well-aligned fibrils with the same polarity, whereas low density or disorder results in a low or even vanishing SHG [86]. In addition, endogenous two-photon excited fluorescence (2PEF) was used to visualize cells in the analyzed tissues. Excitation was provided by a femtosecond titanium–sapphire laser (Mai-Tai, Spectra-Physics) tuned to 860 nm and focused using a water immersion 25× 1.05 NA objective lens (25x, NA 1.05, Plan-Apochromat, Olympus). SHG and 2PEF signals were simultaneously detected in two channels equipped with appropriate spectral filters. Power at the sample was typically 3 to 10 mW. Stacks of 2D images at increasing depth were acquired with 0.34 μm lateral pixel size and 1 μm axial step to generate a 3D data set.

## Statistical analysis

Statistical analysis was performed using Prism 7 (GraphPad Software) with appropriate statistical tests, including the Mann-Whitney test, one-way and two-way ANOVA, or the Kruskal-Wallis test, followed, when necessary, by multiple comparison tests (posthoc analysis). Values of P< 0.05 were considered statistically significant.

## References

[1] Hassell JR, Birk DE. The molecular basis of corneal transparency. Exp Eye Res 2010;91:326–35. 10.1016/j.exer.2010.06.021.

[2] Waring GO, Bourne WM, Edelhauser HF, Kenyon KR. The corneal endothelium. Normal and pathologic structure and function. Ophthalmology 1982;89:531–90.

[3] Zhang J, Sisley AMG, Anderson AJ, Taberner AJ, McGhee CNJ, Patel D V. Characterization of a Novel Collagen Scaffold for Corneal Tissue Engineering. Tissue Eng Part C Methods 2015. 10.1089/ten.TEC.2015.0304.

[4] Lwigale PY, Cressy PA, Bronner-Fraser M. Corneal keratocytes retain neural crest progenitor cell properties. Dev Biol 2005;288:284–93. 10.1016/J.YDBIO.2005.09.046.

[5] Creuzet S, Couly G, Le Douarin NM. Patterning the neural crest derivatives during development of the vertebrate head: insights from avian studies. J Anat 2005;207:447–59. 10.1111/j.1469-7580.2005.00485.x.

[6] Gage PJ, Rhoades W, Prucka SK, Hjalt T. Fate maps of neural crest and mesoderm in the mammalian eye. Invest Ophthalmol Vis Sci 2005;46:4200–8. 10.1167/iovs.05-0691.

[7] Yoshida S, Shimmura S, Shimazaki J, Shinozaki N, Tsubota K. Serum-Free Spheroid Culture of Mouse Corneal Keratocytes. Invest Ophthalmol Vis Sci 2005;46:1653–8. 10.1167/iovs.04-1405.

[8] Davies SB, Di Girolamo N. Corneal stem cells and their origins: significance in developmental biology. Stem Cells Dev 2010;19:1651–62. 10.1089/scd.2010.0201.

[9] Grieve K, Ghoubay D, Georgeon C, Latour G, Nahas A, Plamann K, et al. Stromal striae: A new insight into corneal physiology and mechanics. Sci Rep 2017;7. 10.1038/s41598-017-13194-6.

[10] Maurice DM. THE STRUCTURE AND TRANSPARENCY OF THE CORNEA. vol. 36. 1957.

[11] West-Mays JA, Dwivedi DJ. The keratocyte: corneal stromal cell with variable repair phenotypes. Int J Biochem Cell Biol 2006;38:1625–31. 10.1016/j.biocel.2006.03.010.

[12] Hay ED. Development of the Vertebrate Cornea. In: Bourne GH, Danielli JF, Jeon KW, editors. Int Rev Cytol, vol. 63, Academic Press; 1980, p. 263–322.

[13] Méndez-Maldonado K, Vega-López GA, Aybar MJ, Velasco I. Neurogenesis From Neural Crest Cells: Molecular Mechanisms in the Formation of Cranial Nerves and Ganglia. Front Cell Dev Biol 2020;8:635. 10.3389/FCELL.2020.00635.

[14] Creuzet S, Vincent C, Couly G. Neural crest derivatives in ocular and periocular structures. Int J Dev Biol 2005;49:161–71. 10.1387/ijdb.041937sc.

[15] Lwigale PY. Corneal Development: Different Cells from a Common Progenitor. Prog Mol Biol Transl Sci 2015;134:43–59. 10.1016/BS.PMBTS.2015.04.003.

[16] Du Y, Funderburgh ML, Mann MM, SundarRaj N, Funderburgh JL. Multipotent stem cells in human corneal stroma. Stem Cells 2005;23:1266–75. 10.1634/stemcells.2004-0256.

[17] Meek KM, Knupp C. Corneal structure and transparency. Prog Retin Eye Res 2015;49:1–16. 10.1016/j.preteyeres.2015.07.001.

[18] Takamiya M, Stegmaier J, Kobitski AY, Schott B, Weger BD, Margariti D, et al. Pax6 organizes the anterior eye segment by guiding two distinct neural crest waves. PLoS Genet 2020;16:e1008774. 10.1371/JOURNAL.PGEN.1008774.

[19] Barut Selver MDC and O. Clinical Evaluation of Corneal Neovascularization: A Brief Review. Journal of Ophthalmic Research and Vision Care 2022;2:1–6. 10.54289/JORVC2200106.

[20] Nicholas MP, Mysore N. Corneal neovascularization. Exp Eye Res 2021;202:108363. 10.1016/j.exer.2020.108363.

[21] Loiseau A, Raîche-Marcoux G, Maranda C, Bertrand N, Boisselier E. Animal Models in Eye Research: Focus on Corneal Pathologies. Int J Mol Sci 2023;24:16661. 10.3390/ijms242316661.

[22] Martino VB, Sabljic T, Deschamps P, Green RM, Akula M, Peacock E, et al. Conditional deletion of AP-2β in the cranial neural crest results in anterior segment dysgenesis and early-onset glaucoma. Dis Model Mech 2016. 10.1242/dmm.025262.

[23] Tian H, Sanders E, Reynolds A, van Roy F, van Hengel J. Ocular Anterior Segment Dysgenesis upon Ablation of p120 Catenin in Neural Crest Cells. Investigative Opthalmology & Visual Science 2012;53:5139. 10.1167/iovs.12-9472.

[24] Ting DSJ, Deshmukh R, Ting DSW, Ang M. Big data in corneal diseases and cataract: Current applications and future directions. Front Big Data 2023;6:1017420. 10.3389/FDATA.2023.1017420.

[25] Fujimoto H, Maeda N, Shintani A, Nakagawa T, Fuchihata M, Higashiura R, et al. Quantitative Evaluation of the Natural Progression of Keratoconus Using Three-Dimensional Optical Coherence Tomography. Invest Ophthalmol Vis Sci 2016;57:OCT169–75. 10.1167/IOVS.15-18650.

[26] Swamynathan SK. Ocular surface development and gene expression. J Ophthalmol 2013;2013. 10.1155/2013/103947.

[27] Giovannone D, Ortega B, Reyes M, El-Ghali N, Rabadi M, Sao S, et al. Chicken trunk neural crest migration visualized with HNK1. Acta Histochem 2015;117:255–66. 10.1016/j.acthis.2015.03.002.

[28] Walker H, Taiyab A, Deschamps P, Williams T, West-Mays JA. Conditional deletion of ap-2β in the periocular mesenchyme of mice alters corneal epithelial cell fate and stratification. Int J Mol Sci 2021;22. 10.3390/IJMS22168730.

[29] Jackson BW, Grund C, Schmid E, Bürki K, Franke WW, Illmensee K. Formation of Cytoskeletal Elements During Mouse Embryogenesis: Intermediate Filaments of the Cytokeratin Type and Desmosomes in Preimplantation Embryos. Differentiation 1980;17:161–79. 10.1111/J.1432-0436.1980.TB01093.X.

[30] Merjava S, Brejchova K, Vernon A, Daniels JT, Jirsova K. Cytokeratin 8 Is Expressed in Human Corneoconjunctival Epithelium, Particularly in Limbal Epithelial Cells. Invest Ophthalmol Vis Sci 2011;52:787–94. 10.1167/IOVS.10-5489.

[31] Vanderburg CR, Hay ED. E-cadherin transforms embryonic corneal fibroblasts to stratified epithelium with desmosomes. Acta Anat (Basel) 1996;157:87–104. 10.1159/000147870.

[32] Pontoriero GF, Smith AN, Miller L-AD, Radice GL, West-Mays JA, Lang RA. Co-operative roles for E-cadherin and N-cadherin during lens vesicle separation and lens epithelial cell survival. Dev Biol 2009;326:403–17. 10.1016/j.ydbio.2008.10.011.

[33] Shimamura K, Takeichi M. Local and transient expression of E-cadherin involved in mouse embryonic brain morphogenesis. Development 1992;116:1011–9. 10.1242/dev.116.4.1011.

[34] Gatzioufas Z, Charalambous P, Thanos S. Reduced expression of the gap junction protein Connexin 43 in keratoconus. Eye 2008 22:2 2007;22:294–9. 10.1038/sj.eye.6702972.

[35] Cheung M, Briscoe J. Neural crest development is regulated by the transcription factor Sox9. Development 2003;130:5681–93. 10.1242/DEV.00808.

[36] Menzel-Severing J, Zenkel M, Polisetti N, Sock E, Wegner M, Kruse FE, et al. Transcription factor profiling identifies Sox9 as regulator of proliferation and differentiation in corneal epithelial stem/progenitor cells. Sci Rep 2018. 10.1038/s41598-018-28596-3.

[37] Shibata S, Yasuda A, Renault-Mihara F, Suyama S, Katoh H, Inoue T, et al. Sox10-Venus mice: A new tool for real-time labeling of neural crest lineage cells and oligodendrocytes. Mol Brain 2010;3:1–14. 10.1186/1756-6606-3-31/TABLES/1.

[38] Kuhlbrodt K, Herbarth B, Sock E, Hermans-Borgmeyer I, Wegner M. Sox10, a Novel Transcriptional Modulator in Glial Cells. Journal of Neuroscience 1998;18:237–50. 10.1523/JNEUROSCI.18-01-00237.1998.

[39] Aoki Y, Saint-Germain N, Gyda M, Magner-Fink E, Lee Y-H, Credidio C, et al. Sox10 regulates the development of neural crest-derived melanocytes in Xenopus. Dev Biol 2003;259:19–33. 10.1016/S0012-1606(03)00161-1.

[40] Ma J, Lwigale P. Transformation of the Transcriptomic Profile of Mouse Periocular Mesenchyme During Formation of the Embryonic Cornea. Invest Ophthalmol Vis Sci 2019;60:661–76. 10.1167/iovs.18-26018.

[41] Kreitzer FR, Salomonis N, Sheehan A, Huang M, Park JS, Spindler MJ, et al. A robust method to derive functional neural crest cells from human pluripotent stem cells. Am J Stem Cells 2013;2:119–31.

[42] Martino VB, Sabljic T, Deschamps P, Green RM, Akula M, Peacock E, et al. Conditional deletion of AP-2β in the cranial neural crest results in anterior segment dysgenesis and early-onset glaucoma. Dis Model Mech 2016. 10.1242/dmm.025262.

[43] Carbe C, Garg A, Cai Z, Li H, Powers A, Zhang X. An Allelic Series at the Paired Box Gene 6 (Pax6) Locus Reveals the Functional Specificity of Pax Genes. Journal of Biological Chemistry 2013;288:12130–41. 10.1074/JBC.M112.436865.

[44] Sunny SS, Lachova J, Dupacova N, Kozmik Z. Multiple roles of Pax6 in postnatal cornea development. Dev Biol 2022;491:1–12. 10.1016/J.YDBIO.2022.08.006.

[45] Walther C, Gruss P. Pax-6, a murine paired box gene, is expressed in the developing CNS. Development 1991;113:1435–49. 10.1242/DEV.113.4.1435.

[46] Cvekl A, Tamm ER. Anterior eye development and ocular mesenchyme: new insights from mouse models and human diseases. Bioessays 2004;26:374–86. 10.1002/BIES.20009.

[47] Davis-Silberman N, Kalich T, Oron-Karni V, Marquardt T, Kroeber M, Tamm ER, et al. Genetic dissection of Pax6 dosage requirements in the developing mouse eye. Hum Mol Genet 2005;14:2265–76. 10.1093/HMG/DDI231.

[48] Collinson JM, Morris L, Reid AI, Ramaesh T, Keighren MA, Flockhart JH, et al. Clonal analysis of patterns of growth, stem cell activity, and cell movement during the development and maintenance of the murine corneal epithelium. Developmental Dynamics 2002;224:432–40. 10.1002/dvdy.10124.

[49] Graw J. The genetic and molecular basis of congenital eye defects. Nat Rev Genet 2003;4:876–88. 10.1038/nrg1202.

[50] Xiao H, Hu L, Lin X, Liu L, Dong X, Yang L, et al. Pax6 Directly Regulates the Raver2/sFlt-1 Expression in Corneal Epithelial Cells to Maintain the Cornea’s Avascular Privilege. Invest Ophthalmol Vis Sci 2025;66:52. 10.1167/IOVS.66.4.52.

[51] Espana EM, Birk DE. Composition, structure and function of the corneal stroma. Exp Eye Res 2020;198:108137. 10.1016/j.exer.2020.108137.

[52] Morris SA, Grewal S, Barrios F, Patankar SN, Strauss B, Buttery L, et al. Dynamics of anterior–posterior axis formation in the developing mouse embryo. Nat Commun 2012;3:673. 10.1038/ncomms1671.

[53] Bedzhov I, Bialecka M, Zielinska A, Kosalka J, Antonica F, Thompson AJ, et al. Development of the anterior-posterior axis is a self-organizing process in the absence of maternal cues in the mouse embryo. Cell Res 2015;25:1368–71. 10.1038/cr.2015.104.

[54] Schnell S, Painter KJ, Maini PK, Othmer HG. Spatiotemporal Pattern Formation in Early Development: A Review of Primitive Streak Formation and Somitogenesis, 2001, p. 11–37. 10.1007/978-1-4613-0133-2_2.

[55] van Eeden F, St Johnston D. The polarisation of the anterior-posterior and dorsal-ventral axes during Drosophila oogenesis. Curr Opin Genet Dev 1999;9:396–404. 10.1016/S0959-437X(99)80060-4.

[56] Andoniadou CL, Martinez-Barbera JP. Developmental mechanisms directing early anterior forebrain specification in vertebrates. Cellular and Molecular Life Sciences 2013;70:3739–52. 10.1007/s00018-013-1269-5.

[57] Lo CW, Gilula NB. Gap junctional communication in the preimplantation mouse embryo. Cell 1979;18:399–409. 10.1016/0092-8674(79)90059-X.

[58] Elias LAB, Kriegstein AR. Gap junctions: multifaceted regulators of embryonic cortical development. Trends Neurosci 2008;31:243–50. 10.1016/j.tins.2008.02.007.

[59] Paznekas WA, Boyadjiev SA, Shapiro RE, Daniels O, Wollnik B, Keegan CE, et al. Connexin 43 (GJA1) Mutations Cause the Pleiotropic Phenotype of Oculodentodigital Dysplasia. The American Journal of Human Genetics 2003;72:408–18. 10.1086/346090.

[60] Cella W, Cabral de Vasconcellos JP, Barbosa de Melo M, Kneipp B, Costa FF, Longui CA, et al. Structural Assessment of *PITX2*, *FOXC1*, *CYP1B1*, and *GJA1* Genes in Patients with Axenfeld-Rieger Syndrome with Developmental Glaucoma. Investigative Opthalmology & Visual Science 2006;47:1803. 10.1167/iovs.05-0979.

[61] Tidu A, Ghoubay-Benallaoua D, Teulon C, Asnacios S, Grieve K, Portier F, et al. Highly concentrated collagen solutions leading to transparent scaffolds of controlled three-dimensional organizations for corneal epithelial cell colonization. Biomater Sci 2018;6:1492–502. 10.1039/c7bm01163f.

[62] Sheppard J, Hayes S, Boote C, Votruba M, Meek KM. Changes in Corneal Collagen Architecture during Mouse Postnatal Development. Investigative Opthalmology & Visual Science 2010;51:2936. 10.1167/iovs.09-4612.

[63] Michelacci YM. Collagens and proteoglycans of the corneal extracellular matrix. Brazilian Journal of Medical and Biological Research 2003;36:1037–46. 10.1590/S0100-879X2003000800009.

[64] Sun M, Zafrullah N, Devaux F, Hemmavanh C, Adams S, Ziebarth NM, et al. Collagen XII Is a Regulator of Corneal Stroma Structure and Function. Investigative Opthalmology & Visual Science 2020;61:61. 10.1167/iovs.61.5.61.

[65] Tidu A, Ghoubay-Benallaoua D, Lynch B, Haye B, Illoul C, Allain J-M, et al. Development of human corneal epithelium on organized fibrillated transparent collagen matrices synthesized at high concentration. Acta Biomater 2015;22. 10.1016/j.actbio.2015.04.018.

[66] Daxer A, Fratzl P. Collagen fibril orientation in the human corneal stroma and its implication in keratoconus. Invest Ophthalmol Vis Sci 1997;38:121–9.

[67] Norouzpour A, Mehdizadeh A. A Novel Insight into Keratoconus: Mechanical Fatigue of the Cornea. Medical Hypothesis, Discovery and Innovation in Ophthalmology 2012;1:14–7.

[68] Ghoubay D, Borderie M, Grieve K, Martos R, Bocheux R, Nguyen T-M, et al. Corneal stromal stem cells restore transparency after N2 injury in mice. Stem Cells Transl Med 2020;9. 10.1002/sctm.19-0306.

[69] Coulombre AJ, Coulombre JL. Corneal development: III. The role of the thyroid in dehydration and the development of transparency. Exp Eye Res 1964;3:105–14. 10.1016/S0014-4835(64)80024-5.

[70] Babushkina A, Lwigale P. Periocular neural crest cell differentiation into corneal endothelium is influenced by signals in the nascent corneal environment. Dev Biol 2020;465:119–29. 10.1016/j.ydbio.2020.06.012.

[71] Cogan DG, Kinsey VE. Physiologic Studies on the Cornea. Science (1979) 1942;95:607–8. 10.1126/science.95.2476.607.

[72] Cox JL, Farrell RA, Hart RW, Langham ME. The transparency of the mammalian cornea. J Physiol 1970;210:601–616.4.

[73] Hart RW, Farrell RA. Light scattering in the cornea. J Opt Soc Am 1969;59:766–74. 10.1364/JOSA.59.000766.

[74] Benedek GB. Theory of Transparency of the Eye. Appl Opt 1971;10:459. 10.1364/AO.10.000459.

[75] Twersky V. Transparency of pair-correlated, random distributions of small scatterers, with applications to the cornea. J Opt Soc Am 1975;65:524–30. 10.1364/JOSA.65.000524.

[76] Meek KM, Boote C. The use of X-ray scattering techniques to quantify the orientation and distribution of collagen in the corneal stroma. Prog Retin Eye Res 2009;28:369–92. 10.1016/J.PRETEYERES.2009.06.005.

[77] Koudouna E, Winkler M, Mikula E, Juhasz T, Brown DJ, Jester J V. Evolution of the vertebrate corneal stroma. Prog Retin Eye Res 2018;64:65–76. 10.1016/j.preteyeres.2018.01.002.

[78] Dubois A, Grieve K, Moneron G, Lecaque R, Vabre L, Boccara C. Ultrahigh-resolution full-field optical coherence tomography. Appl Opt 2004;43:2874–83. 10.1364/ao.43.002874.

[79] Grieve K, Paques M, Dubois A, Sahel J, Boccara C, Le Gargasson J-F. Ocular tissue imaging using ultrahigh-resolution, full-field optical coherence tomography. Invest Ophthalmol Vis Sci 2004;45:4126–31. 10.1167/iovs.04-0584.

[80] Ghouali W, Grieve K, Bellefqih S, Sandali O, Harms F, Laroche L, et al. Full-field optical coherence tomography of human donor and pathological corneas. Curr Eye Res 2015;40:526–34. 10.3109/02713683.2014.935444.

[81] Dubois A, Levecq O, Azimani H, Davis A, Ogien J, Siret D, et al. Line-field confocal time-domain optical coherence tomography with dynamic focusing. Opt Express 2018;26:33534. 10.1364/OE.26.033534.

[82] Ogien J, Levecq O, Azimani H, Dubois A. Dual-mode line-field confocal optical coherence tomography for ultrahigh-resolution vertical and horizontal section imaging of human skin in vivo. Biomed Opt Express 2020;11:1327. 10.1364/BOE.385303.

[83] Renier N, Adams EL, Kirst C, Wu Z, Azevedo R, Kohl J, et al. Mapping of Brain Activity by Automated Volume Analysis of Immediate Early Genes. Cell 2016;165:1789–802. 10.1016/j.cell.2016.05.007.

[84] Raoux C, Chessel A, Mahou P, Latour G, Schanne-Klein M-C. Unveiling the lamellar structure of the human cornea over its full thickness using polarization-resolved SHG microscopy. Light Sci Appl 2023;12:190. 10.1038/s41377-023-01224-0.

[85] Raoux C, Schmeltz M, Bied M, Alnawaiseh M, Hansen U, Latour G, et al. Quantitative structural imaging of keratoconic corneas using polarization-resolved SHG microscopy. Biomed Opt Express 2021;12:4163–78. 10.1364/BOE.426145.

[86] Bancelin S, Aimé C, Gusachenko I, Kowalczuk L, Latour G, Coradin T, et al. Determination of collagen fibril size via absolute measurements of second-harmonic generation signals. Nat Commun 2014;5:4920. 10.1038/ncomms5920.

[87] Chen X, Nadiarynkh O, Plotnikov S, Campagnola PJ. Second harmonic generation microscopy for quantitative analysis of collagen fibrillar structure. Nat Protoc 2012;7:654–69. 10.1038/NPROT.2012.009.

