## Supplemental figures for "Role of Neural Crest Cells in Establishing Corneal Transparency During Embryonic Development in Mice"

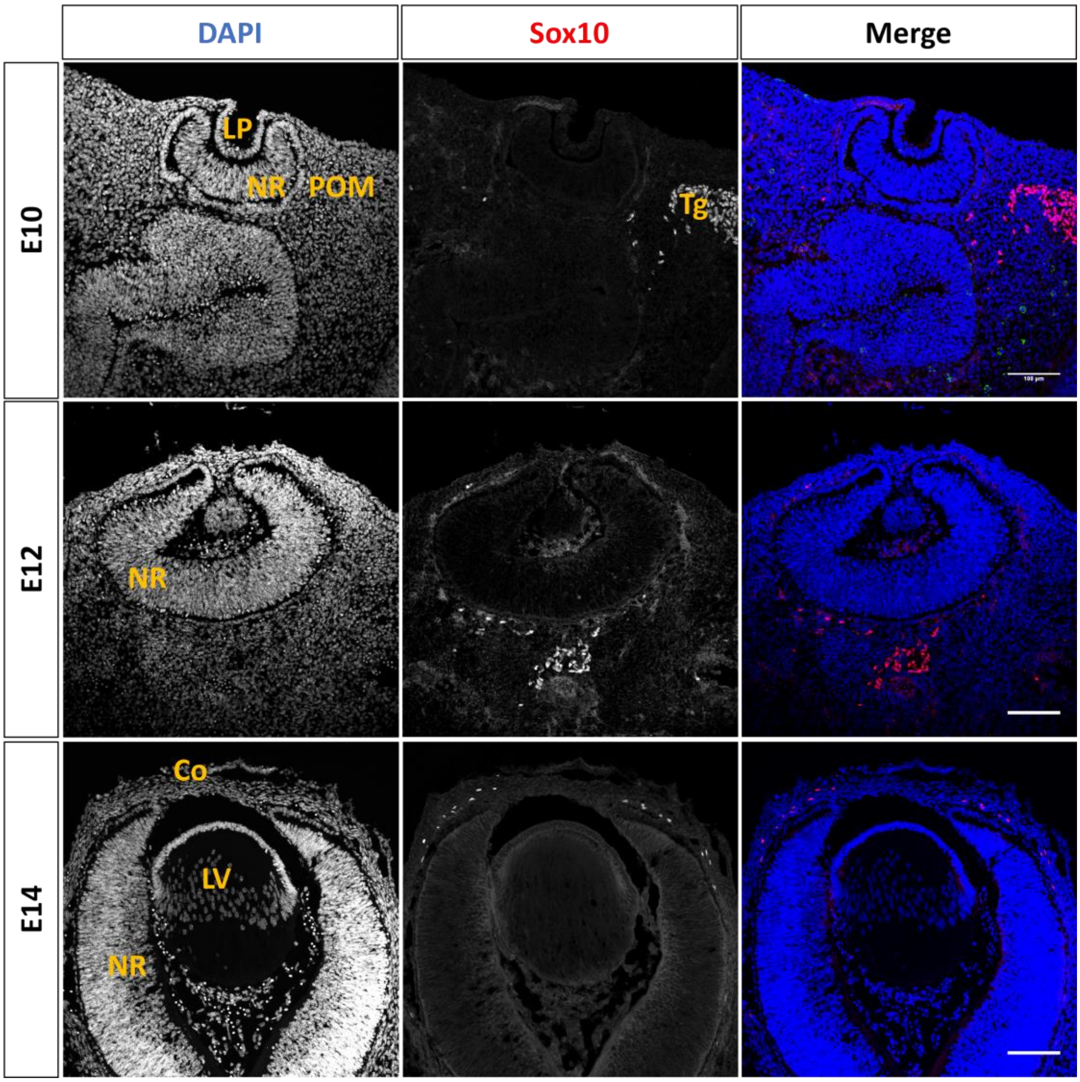


**S1 Figure. Sox 10 immunostaining in mouse embryos cross sections**

Immunostaining images of E10, E12, and E14 showing wider views of the optic vesicle, complementing Figureure 3, to better visualize the Sox10-positive cells.

Co, Cornea; LP, Lens Pit; LV, lens vesicle; NR, neuroretina; POM, Periocular mesenchyme. Scale bar=100µm

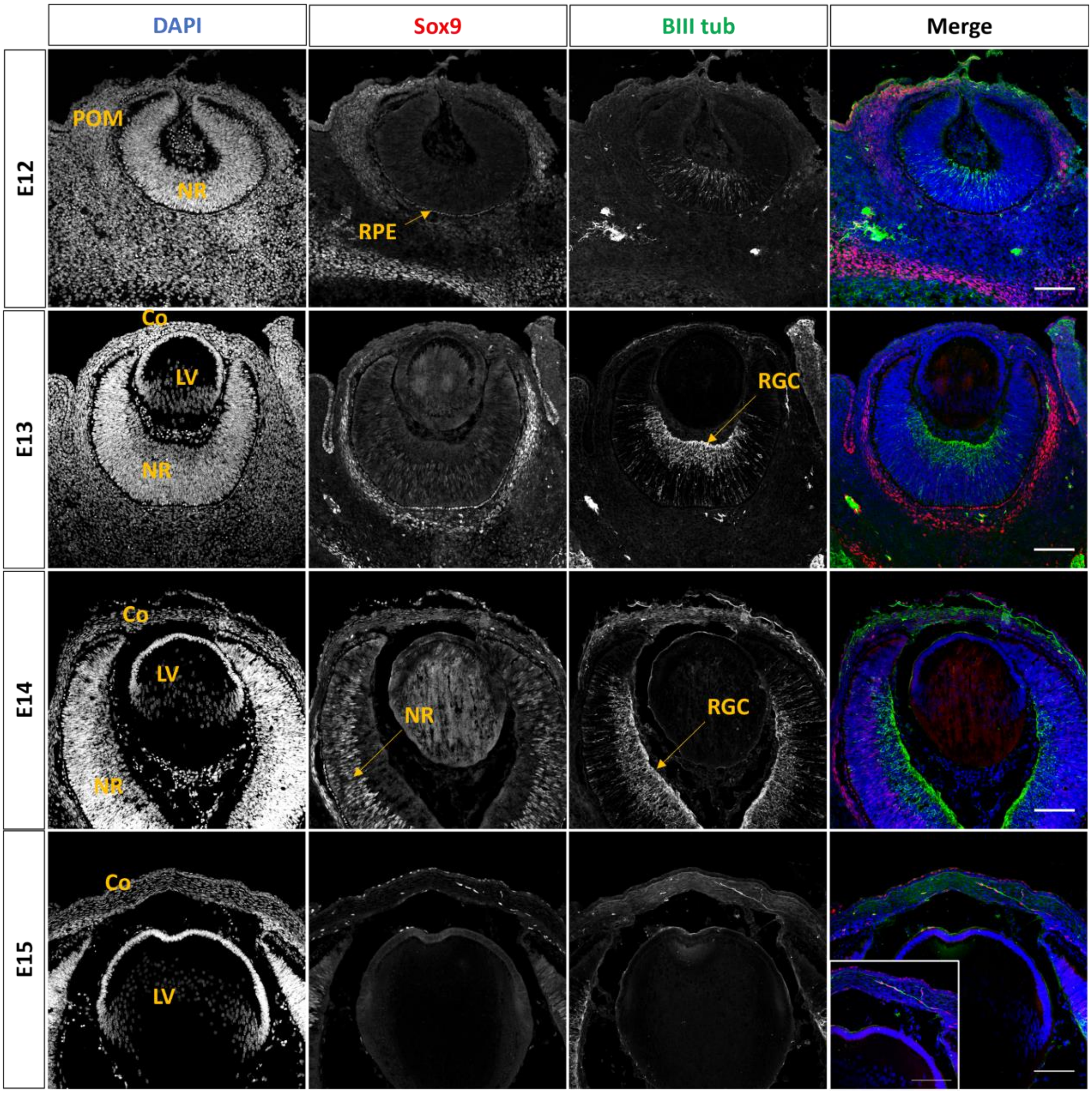


**S2 Figure. Sox9 and βIII tubulin immunostaining in embryos cross sections**

Images from E12 to E15 showing wider views to visualize the co-localization of Sox9 and βIII tubulin in the embryonic eyes.

Co, Cornea; LV, lens vesicle; NR, neuroretina; RPE, Retinal Pigmented Epithelium; RGC, Retinal Ganglion Cell; POM, Periocular mesenchyme. Scale bar=100µm.

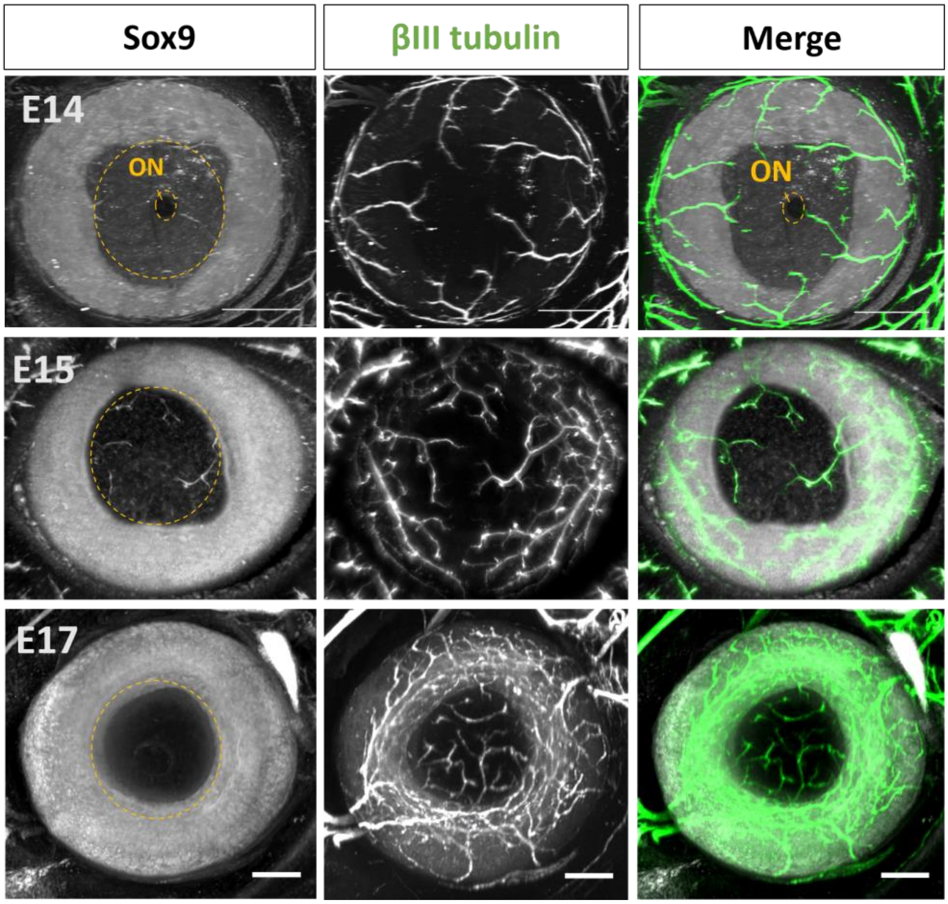


**S3 Figure. Innervation process intensifying as development progresses**

Frontal view of the eye at E14, E15 and E17 showing Sox9, βIII tubulin, and the merge staining of the two markers. ON: Optic nerve. Scale bar: 100µm

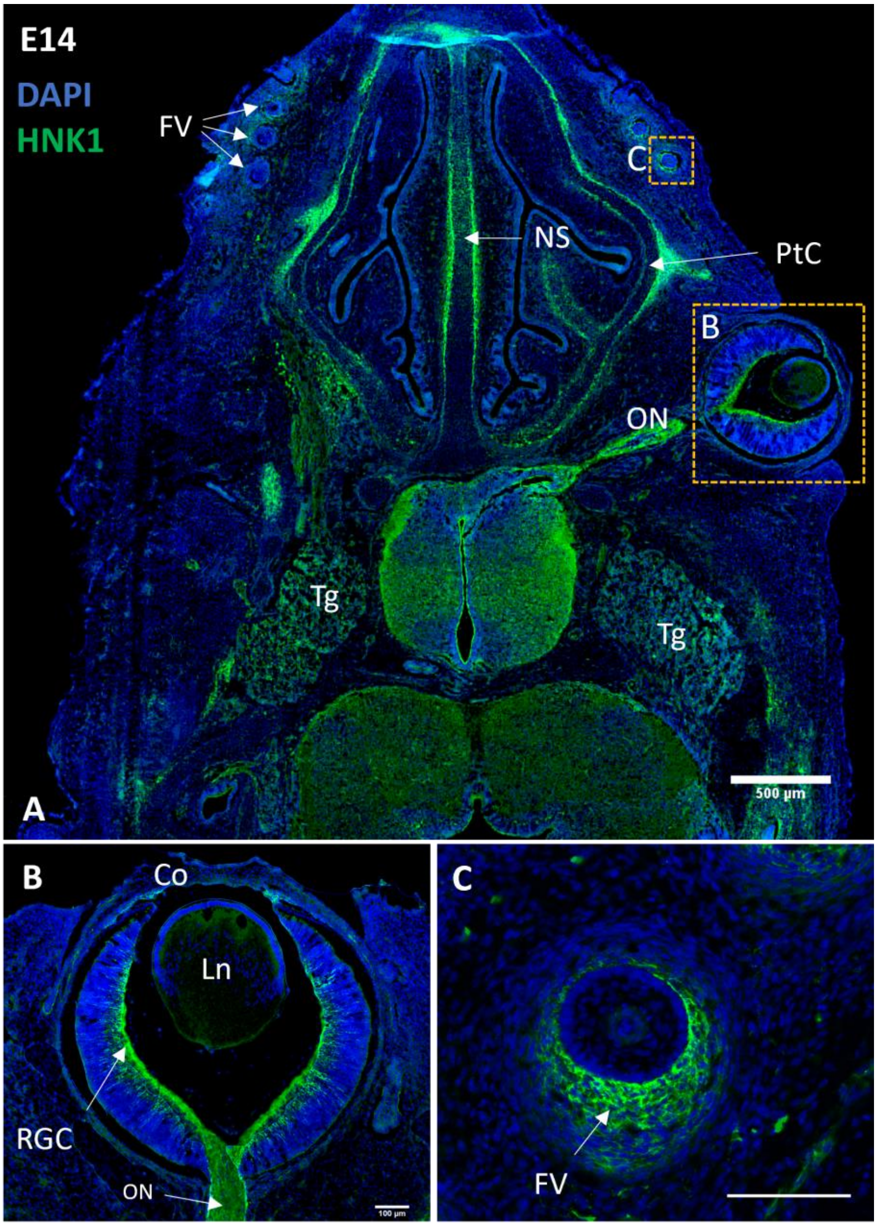


**S4 Figure. HNK1 staining in different region of the head**

HNK1, is widely expressed in tissues associated with neural crest cell derivatives.

CO, cornea; FV, Follicles of vibrissae; NS, Nasal Septum; ON: Optic nerve; PtC, Parietotectal Cartilage; RGC, Retinal ganglion cell; TG: Trigeminal Ganglion.

Scale bar, A: 500µm, B and C: 100µm.

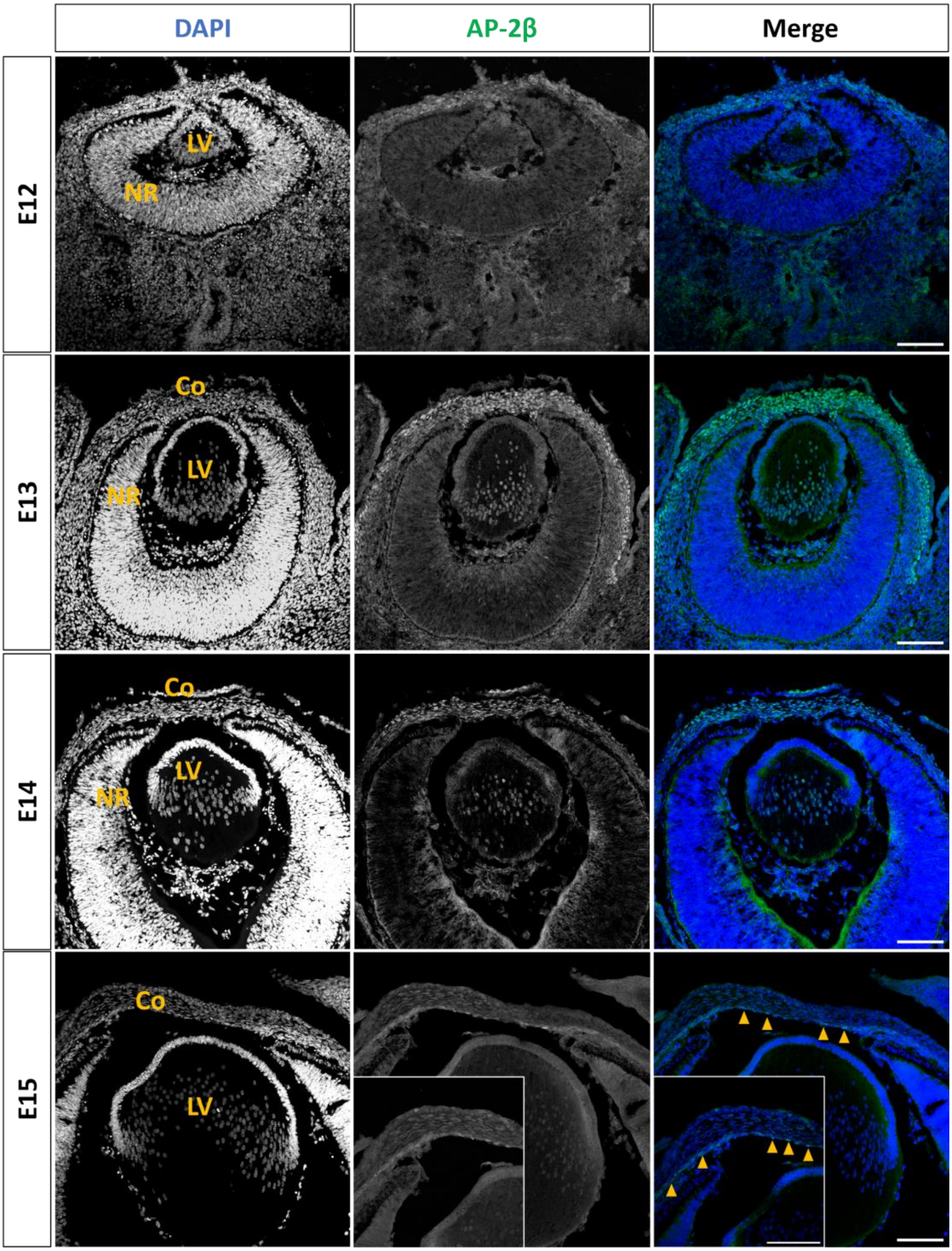


**S5 Figure. AP2β immunostaining in mouse embryos cross sections**

Images from E12 to E15 showing wider views to visualize AP-2β in the embryonic eyes.

Co, Cornea; LV, lens vesicle; NR, neuroretina; Arrow head show the endothelial layer. Scale bar=100µm.

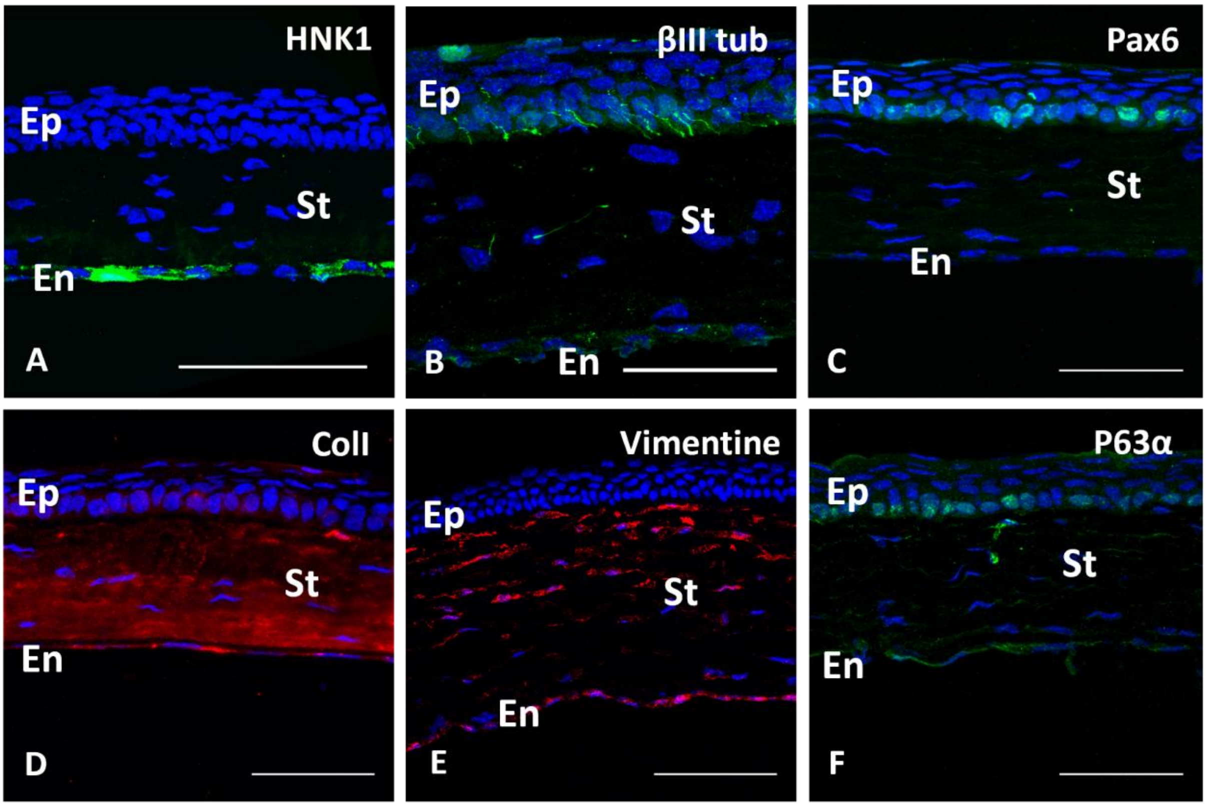


**S6 Figure. Adult mouse cornea phenotyping**

**(A-F)** Immunostaining of adult mouse cornea cross sections**.** Some endothelial cells stained with HNK1**,** nerves by BIII tub (B). Identification of keratocytes with vimentin staining in the stromal layer (E), epithelial stem cells (Pax6 and p63 alpha) in the basal epithelial layer (C and F) and Collagen I in the stromal layer (D). Scale bar: 100µm

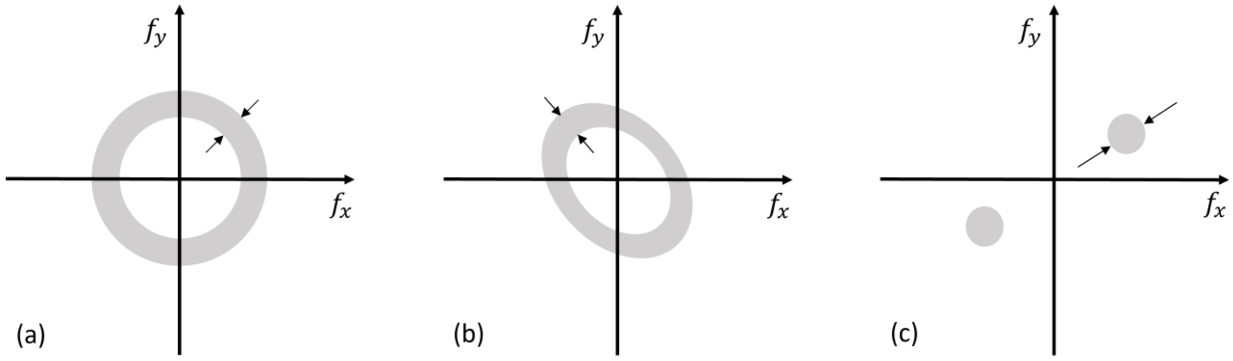


**S7 Figure Spectral measurement of interfibrillar distance**

If the cut performed for TEM imaging is perpendicular to the fibril structure, this gives rise to an annular structure of the Fourier spectrum, the radius of which is inversely related to the interfibrillar distance (a). The upper and lower limit for the radius was estimated from the FFT window. If the fibrils are cut with an angle different from 90°, these values were determined on the long axis of the elliptical Fourier spectrum (b). If the fibrils lie in the plane of the cut, two distinct spots appear in the Fourier spectrum, which again permits the determination of the upper and lower limit (c). Given that the interfibril distance is inversely related to distances in frequency space, the upper limit obtained for the interfibrillar distance may be slightly overestimated. It is essential to have homogeneous regions where the fibers are clearly distinguishable.

### Table S1. Reagent and resources used in the study

| REAGENT or RESOURCE | SOURCE | IDENTIFIER |
|---|---|---|
| **Antibodies** | | |
| **AP-2β** | Cell Signaling | 2509 |
| **E-Cadherin** | R&D Systems | AF748 |
| **CK15** | Abcam | ab52816 |
| **CK18** | Abcam | ab93741 |
| **Collagen I** | Abcam | ab6308 |
| **Collagen IV** | Abcam | Ab6586 |
| **Collagen V** | Abcam | ab7046 |
| **Connexin 43** | Santa Cruz | sc-271837 |
| **HNKI** | Sigma-Aldrich | C6680 |
| **Ki67** | Abcam | ab15580 |
| **P63α** | Cell Signaling | 4892S |
| **PAX6** | Millipore | AB2237 |
| **Sox9** | R&D Systems | AF3075 |
| **Sox10** | Santa Cruz | sc-17342 |
| **βIII-Tubulin** | Sigma | T2200 |
| **Vimentin** | Abcam | ab92547 |
| **Anti-Mouse, Alexa Fluor 488** | Sigma | SAB4600029 |
| **Anti-Mouse, Alexa Fluor 594** | Sigma | SAB4600092 |
| **Anti-Goat, Alexa Fluor 488** | Sigma | SAB4600028 |
| **Anti-Goat, Alexa Fluor 594** | Sigma | SAB4600091 |
| **Anti-Rabbit, Alexa Fluor 488** | Sigma | SAB4600030 |
| **Anti-Rabbit, Alexa Fluor 594** | Sigma | SAB4600093 |
| **Anti-Rabbit, Alexa Fluor 790** | Abcam | Ab186693 |
| **Chemicals** | | |
| **PFA** | Sigma | 178127 |
| **PBS** | Life Technologie | 14190185 |
| **Thimerosal** | Sigma | T5125-10 |
| **Sucrose** | Sigma | S7903 |
| **Gelatin** | Sigma | G1393 |
| **Isopentane** | VWR | 24872-298 |
| **Triton X100** | Sigma | T8787 |
| **Hematoxylin** | 15/20 hospital | |
| **Ponceau** | 15/20 hospital | |
| **Fuchsin** | 15/20 hospital | |
| **Phosphomolybdic acid** | 15/20 hospital | |
| **Acetic acid** | VWR | 201003-295 |
| **Eukitt** | Sigma | 3989 |
| **Ethanol** | VWR | 20821-365 |
| **Agarose** | Merck | A2576 |
| **Methanol** | VWR | 20847-360 |
| **DCM** | Sigma | 5-89581 |
| **DBE** | Sigma | 108014 |
| **Fluoromount G** | Cliniscience | 0100-01 |
